# Enhanced Sequence-Activity Mapping and Evolution of Artificial Metalloenzymes by Active Learning

**DOI:** 10.1101/2024.02.06.579157

**Authors:** Tobias Vornholt, Mojmír Mutný, Gregor W. Schmidt, Christian Schellhaas, Ryo Tachibana, Sven Panke, Thomas R. Ward, Andreas Krause, Markus Jeschek

## Abstract

Tailored enzymes hold great potential to accelerate the transition to a sustainable bioeconomy. Yet, enzyme engineering remains challenging as it relies largely on serendipity and is, therefore, highly laborious and prone to failure. The efficiency and success rates of engineering campaigns may be improved substantially by applying machine learning to construct a comprehensive representation of the sequence-activity landscape from small sets of experimental data. However, it often proves challenging to reliably model a large protein sequence space while keeping the experimental effort tractable. To address this challenge, we present an integrated pipeline combining large-scale screening with active machine learning and model-guided library design. We applied this strategy to efficiently engineer an artificial metalloenzyme (ArM) catalysing a new-to-nature hydroamination reaction. By combining lab automation and next-generation sequencing, we acquired sequence-activity data for several thousand ArM variants. We then used Gaussian process regression to model the activity landscape and guide further screening rounds according to user-defined objectives. Crucial characteristics of our enhanced enzyme engineering pipeline include i) the cost-effective generation of information-rich experimental data sets, ii) the integration of an explorative round to improve the performance of the model, as well as iii) the consideration of experimental noise during modelling. Our approach led to an order-of-magnitude boost in the hit rate of screening while making efficient use of experimental resources. Smart search strategies like this should find broad utility in enzyme engineering and accelerate the development of novel biocatalysts.

## Introduction

Biocatalysis and metabolic engineering offer sustainable production routes for many compounds of interest and thus hold the potential to transform various industries. However, extensive enzyme engineering is typically required to obtain a suitable biocatalyst for a desired application. This is often a time-consuming, empirical process whose outcome is subject to chance, as classical methods are agnostic to the topology of the underlying sequence-activity landscape. Engineering strategies that incorporate machine learning to model this landscape could render enzyme engineering more efficient and increase the likelihood of identifying an optimal solution. Accordingly, machine learning-assisted directed evolution (MLDE) has attracted significant attention in recent years^1–3^.

In general, MLDE starts with an initial screening round in which both sequence and activity are recorded for a number of enzyme variants. These sequence-activity data are then used to train a machine learning model, with the objective of predicting the activity of untested variants directly from their sequence. If successful, such models can suggest variants that are likely to be highly active and thus support further screening rounds by *in silico* library design^1^. Further, the model can be iteratively updated with new data to improve its predictive performance, a strategy referred to as active learning. While several studies have demonstrated the general feasibility of such approaches^4–12^, there are still various challenges that need to be addressed to maximize the success rate and efficiency of MLDE and enable its widespread implementation. This pertains to various aspects such as library design, experimental data acquisition, model development, and the strategy for sampling the sequence space.

With regard to library design, the crucial challenge is to create a library that is as information-dense as possible to allow for the development of accurate models while keeping the screening effort manageable. In the initial stages of model development, this calls for libraries that exhibit a high degree of sequence diversity to provide adequate information on the underlying sequence space, while at the same time containing a sufficient number of active mutants^13^. These requirements can be difficult to reconcile, as simultaneous randomization of multiple residues commonly results in a large fraction of inactive mutants, from which little to no meaningful information for model training can be extracted.

Once a library has been generated, it is often challenging to measure a sufficiently large set of sequence-activity data. In some cases, high-throughput assays such as fluorescence-activated cell sorting can be combined with deep sequencing to obtain very large data sets^14,15^. However, most enzymatic reactions of industrial relevance require more laborious analytical procedures to obtain a readout for activity. Moreover, the need to also obtain sequence information on all tested variants can lead to prohibitive costs if conventional Sanger sequencing is used. Consequently, most studies to date have relied on small data sets (10^1^-10^2^ variants)^4–10^. While this has led to several successful demonstrations of MLDE, larger data sets are likely to lead to more accurate machine learning models and improve the chances of identifying variants with the desired properties^11^, particularly as the search space increases in size.

Beyond these experimental considerations, several critical decisions have to be made regarding the machine learning strategy. Prominent examples in this regard include the encoding strategy for the protein sequences and the choice of a suitable machine learning algorithm. Many encoding strategies have been suggested for creating a meaningful representation of protein variants, ranging from simple one-hot encoding and descriptors based on amino acid properties^16–18^ to structure-based descriptors^19,20^ and learned embeddings^21,22^. Similarly, various machine learning algorithms have been employed or suggested for MLDE, including linear regression^23–25^, Gaussian processes^4,7–9,25,26^, and neural networks^12^. While the best strategy depends on the data set and task at hand, Gaussian processes have repeatedly revealed their utility for active learning^8,9,25^.

Less attention has been devoted to other aspects of the machine learning process, such as the handling of experimental noise or the sampling strategy during ML-guided screening rounds, both of which are critical to the success and efficiency of MLDE. With regard to the sampling strategy, many studies have relied on a single training phase followed by greedy sampling of the top predictions of the resulting model. Due to inevitable biases in library generation and the limitations in generating sufficient sequence-activity data, this is unlikely to result in a comprehensive and accurate representation of the sequence-activity landscape. Consequently, such models may be “blind” for promising regions of the sequence space, leading to suboptimal outcomes such as low hit rates. Active learning strategies that improve the model in iterative cycles of experiments and machine learning may help to develop a better representation of the sequence-activity landscape, as these can converge to the optimal solution over time^27^. However, the aforementioned bottleneck in experimental data generation makes performing many iterations undesirable. Thus, resources invested into model improvement (i.e., exploration) must be carefully weighed against the focus on regions of the sequence space that are likely to contain active variants but might only comprise local optima (exploitation). In addition, activity may not be the only selection criterion during exploitation. Instead, it is often desirable to sample various potential optima to obtain a diverse set of variants, which requires more elaborate approaches than simple greedy selection of top predictions^28^. Hence, smart sampling strategies for active learning are required to maximize the chances of success at a given experimental budget.

In this study, we introduce an integrated experimental and computational pipeline that addresses critical limitations in the MLDE of enzymes. Specifically, we combine informed library design with large-scale screening and a novel active machine-learning strategy. As an impactful testbed, we selected an artificial metalloenzyme (ArM) for gold-catalysed hydroamination, a new-to-nature reaction for atom-economical C-N bond formation. We simultaneously engineered five crucial amino acid residues in this ArM, corresponding to a search space of 3,200,000 possible variants. To sample this space, we combined lab automation with a cost-efficient next-generation sequencing (NGS) strategy, which allowed us to acquire sequence-activity data on more than 2,000 ArM variants. Furthermore, we developed a machine learning model based on Gaussian process regression that incorporates optimized descriptors and estimates of experimental noise to efficiently navigate the sequence space. Guided by the model’s uncertainty estimates, we performed a second screening round focused on exploration and model refinement. Importantly, our results demonstrate that this targeted exploration substantially improved the model’s performance. The optimized model reliably proposed highly active ArM variants in a final exploitation round, as illustrated by a 12-fold increased hit rate compared to the initial library.

## Results

### Design of an information-dense ArM library

ArMs are hybrid catalysts that promise to significantly increase the number of reactions available in biocatalysis by equipping enzymes with the catalytic versatility of abiotic transition metal cofactors^29^. ArMs have been created for a variety of natural and non-natural reactions^30–35^, and some have demonstrated catalytic prowess comparable to that of natural enzymes^36–39^. However, most ArMs initially display a low activity, and extensive protein engineering is required to identify catalytically proficient variants. This engineering is typically a labour-intensive and slow process. Therefore, ArMs represent an impactful yet challenging use case for MLDE.

A particularly versatile strategy for creating ArMs is to incorporate an organometallic cofactor into the tetrameric protein streptavidin (Sav) using a biotin moiety as the anchor. Using this approach, we have previously engineered an ArM for gold-catalysed hydroamination by exhaustively screening a library of 400 Sav double mutants (Sav S112X K121X) using a whole-cell assay in 96-well plates^40^. While this represents an attractive starting point, extending the search space to more positions offers the opportunity to achieve further improvements, which will be crucial for adapting ArMs for real-world applications. However, exhaustive screening quickly becomes intractable in this case, and smart heuristics for the efficient exploration of the underlying sequence-activity landscape are essential^41^.

To navigate the sequence-activity landscape of the ArM, we devised an iterative active learning cycle involving library design, cloning, screening, and machine learning (Fig. 1a). With regard to library design, the first step is to choose the target residues and a randomization scheme. To maximize the potential impact of the screening campaign, we aimed to find important positions in Sav besides the previously identified residues S112 and K121^40^. Thus, we individually randomized the 20 residues closest to the biotinylated gold cofactor in Sav S112F K121Q, which is the most active variant we had observed before^40^ (referred to as “reference variant” herein). Randomization was performed using degenerate NDT (N = A, C, G or T; D = A, G or T) codons, which encode 12 amino acids covering all chemical classes of amino acids, a strategy that has revealed high success rates at a reduced screening effort^40^. Subsequently, we measured hydroamination activity using our previously established protocol relying on periplasmic catalysis in *Escherichia coli* (Fig. 1b)^40^. We tested 36 clones per randomized position to achieve a statistical library coverage of approximately 95 %^42^. As expected, most variants displayed reduced activity compared to the reference variant (Fig. 1c). Notably, positions 111, 118, and 119 revealed the highest potential for improvement upon mutagenesis, with several variants outperforming the reference variant. Consequently, we selected these positions for further engineering. In addition, we chose to also randomize positions 112 and 121 again, as our observations had indicated that epistatic effects play an important role in highly active ArM mutants^40^.

**Fig. 1.**
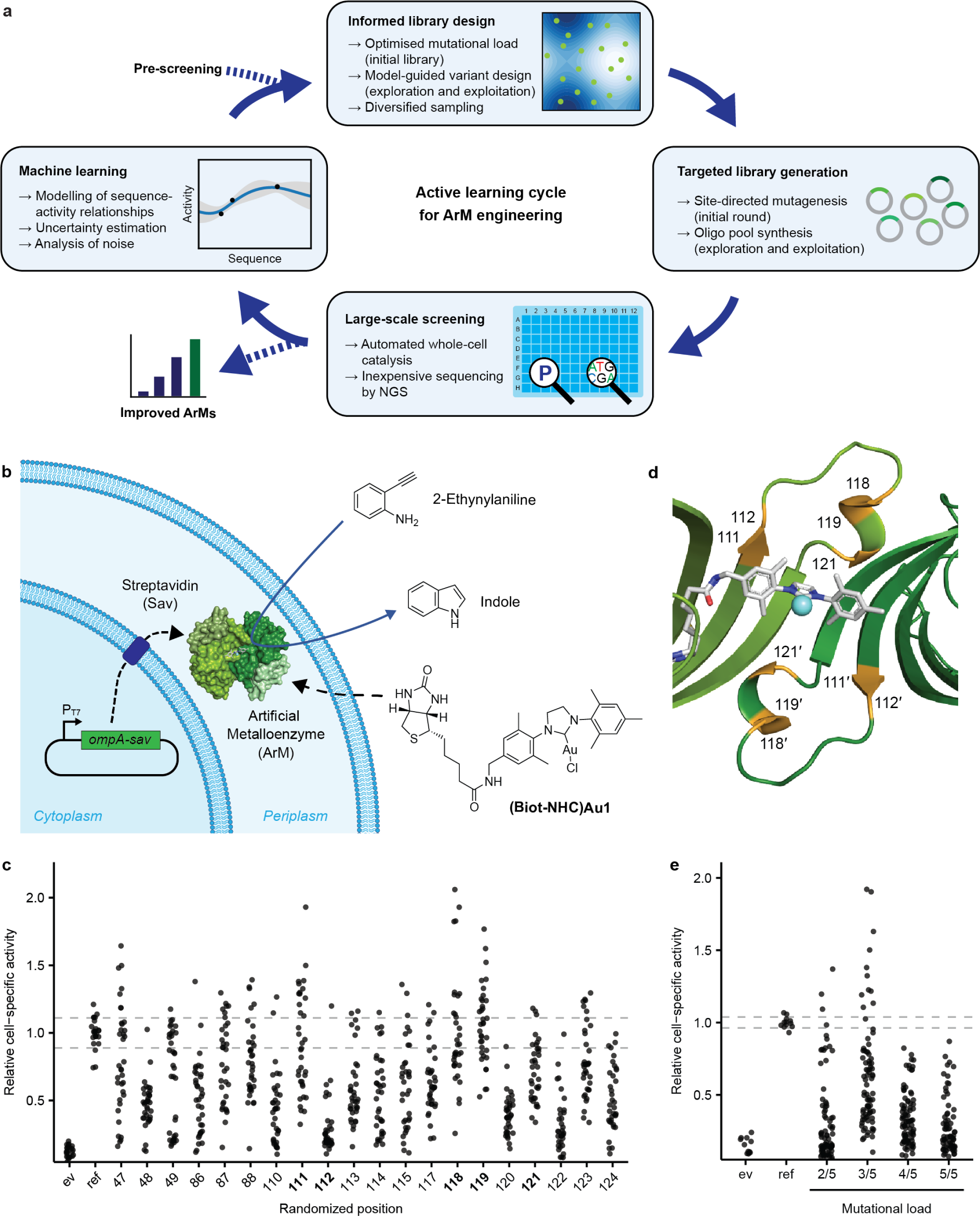
Engineering strategy and library design for ArMs catalysing hydroamination. **a,** Illustration of the active learning strategy for ArM engineering. An iterative process of library design, cloning, large-scale screening and machine learning was used to model the sequence-activity landscape and identify improved ArMs. Crucial steps and considerations are highlighted and are explained in the main text. **b**, Illustration of whole-cell biocatalysis using an ArM in the periplasm of *E. coli*. Sav is exported to the periplasm by means of an N-terminal OmpA signal peptide, where it binds the biotinylated cofactor **(Biot-NHC)Au1**. The resulting ArM converts 2-ethynylaniline to indole in a new-to-nature hydroamination reaction. Indole can subsequently be quantified using a colorimetric assay. **c**, Single site-saturation mutagenesis to identify influential amino acid residues with respect to ArM activity. Starting from the reference variant Sav S112F K121Q, 20 residues in Sav were individually mutated using degenerate NDT codons. The activity of the resulting variants is displayed relative to the mean activity of the reference variant (“ref”). Dashed lines indicate one standard deviation around the mean activity of the reference variant, which was measured in triplicate in each 96-well plate. A strain lacking Sav, i.e., containing an empty vector (“ev”), was included as a control (n = 3 per 96-well plate). The five positions selected for combinatorial randomization are highlighted in bold. Note that no improvement was expected at positions 112 and 121, as the reference variant had already been optimized with regard to these positions^40^. **d**, Residues selected for randomization (highlighted in orange) in a ribbon model of Sav harbouring a metathesis catalyst (PDB 5IRA). For clarity, only two biotin-binding sites of two opposing Sav monomers (a so-called functional dimer) are displayed. **e**, Effect of different multi-site randomization strategies on the activity distribution of ArM libraries. Starting from the reference variant, either two, three, four or five residues amongst positions 111, 112, 118, 119, and 121 were randomized simultaneously. Hydroamination activity is displayed relative to the average activity of the reference variant (“ref“, n = 3 per 96-well plate) for 90 variants from each library. A strain containing an empty vector (“ev”) was included as a control (n = 3 per 96-well plate).

Next, we sought to create a combinatorial library of the five selected positions (111, 112, 118, 119, and 121, Fig. 1d), which, upon full randomization, corresponds to a search space of 20^5^ = 3,200,000 variants. This greatly exceeds the capacity of typical activity assays and well plate-based screenings. Thus, navigating the underlying sequence-activity landscape represents a significant challenge. In order to model this space for MLDE, it is crucial to design a library that offers a good coverage of the targeted sequence space and at the same time maintains a sufficient proportion of active variants^13^. While simultaneous randomization of all five residues would fulfil the first criterion, we anticipated that the high mutational load would likely lead to a large fraction of inactive variants. This would not only diminish the chances of identifying improved variants but, importantly, would be uninformative for machine learning. Upon initial tests, we indeed observed a marked drop in the activity distribution when randomizing more than three of the five positions simultaneously (Fig. 1e). Accordingly, we set out to construct a library with three to four mutations distributed across the five target residues as a good compromise between high sequence-diversity and sufficient residual activity. In other words, the constructed library covers all five target positions, but individual variants contain at most four amino acid substitutions relative to the reference variant Sav S112F K121Q, which served as the parent of the library (Supplementary Fig. 1). This was achieved by site-directed mutagenesis PCR using various sets of primers containing degenerate NNK (K = G or T) codons at different positions and subsequent mixing of the resulting sub-libraries (see Methods).

### Large-scale acquisition of sequence-activity data

Our previously established whole-cell screening protocol for ArMs relied on periplasmic Sav expression, ArM assembly and catalysis in 96-well plate format. By combining this protocol with conventional Sanger sequencing, we were able to obtain sequence-activity data for a few hundred variants^40^. Although this platform was more flexible and simpler than comparable screening strategies involving protein purification, it still required considerable manual labour, particularly for product quantification. Additionally, when larger data sets are required, Sanger sequencing rapidly leads to prohibitively high sequencing costs. To facilitate the generation of larger data sets for MLDE, we thus sought to minimize manual intervention in the activity assay and develop more cost-efficient means of obtaining the sequence information for each functionally characterised variant.

First, we automated all steps in the assay protocol that are labour-intensive (and thus limiting in terms of throughput) or critical for reproducibility. Specifically, we made use of a Tecan EVO 200 platform for all steps from colony picking to product quantification, with the exception of Sav expression in 96-deep well plates, which only requires a small number of pipetting steps (Fig. 2a). The most important addition to our previous semi-automated pipeline^40^ is the photometric quantification of the product indole. While this is a laborious procedure when carried out manually, the automated version simplifies screenings and proved to be very reproducible (Supplementary Fig. 2). As the robotic platform can handle up to eight 96-well plates at the same time, it greatly accelerates the acquisition of large data sets.

**Fig. 2.**
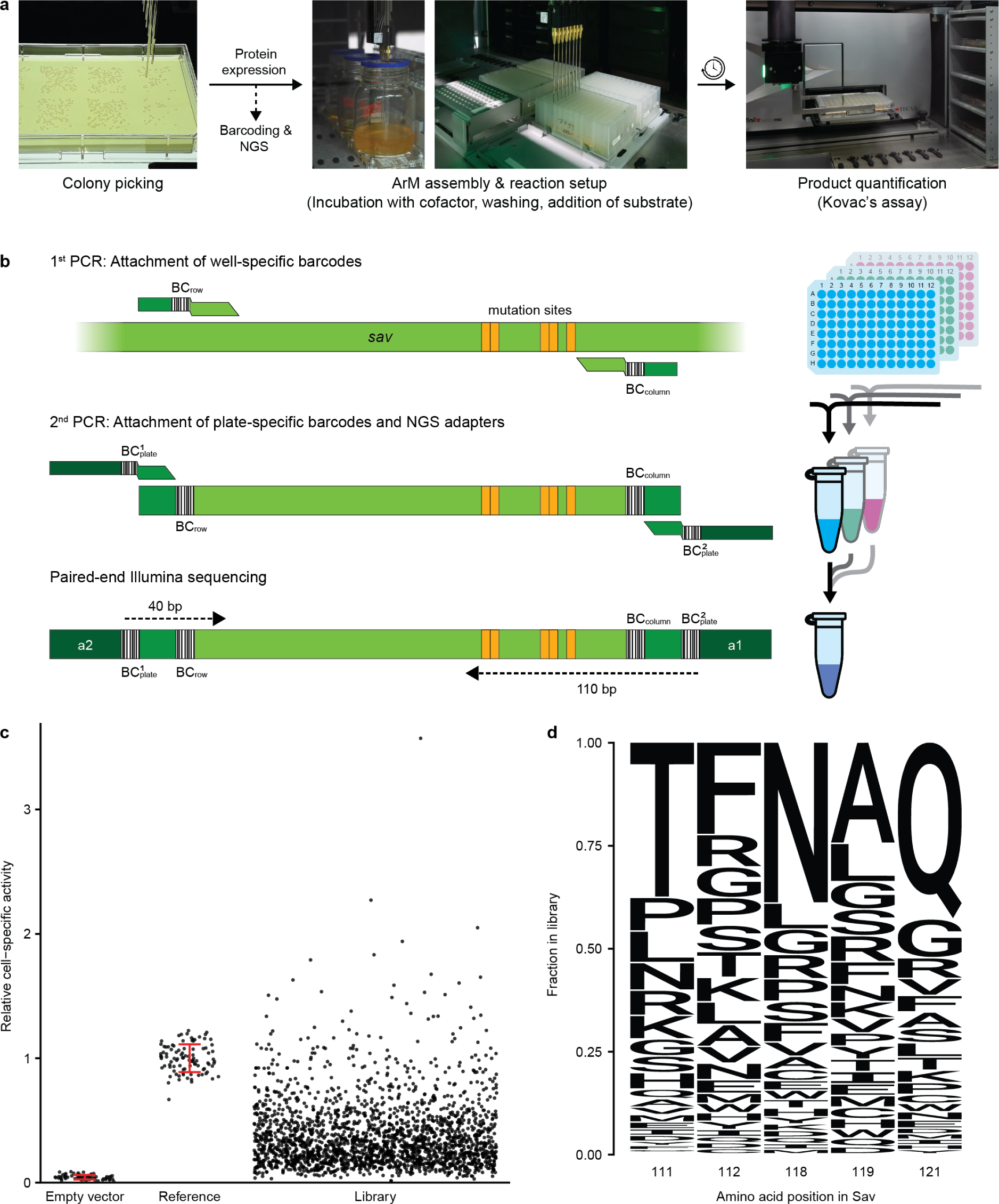
Large-scale acquisition of sequence-activity data for ArMs. **a,** Depiction of the critical automated steps in the screening workflow. Colony picking, ArM assembly, reaction setup, and product quantification were performed on a lab automation platform. The less labour-intensive protein expression protocol was performed manually. In parallel to the activity assay, samples of the starter cultures were processed further for NGS. **b**, PCR-based barcoding strategy for cost-effective sequencing of Sav variants in 96-well plates by NGS. First, the mutated region of the Sav gene is amplified using primers with row- (BC_row_) and column-specific (BC_column_) DNA barcodes. This step is performed in PCR plates using heat-treated bacterial cultures as templates. After pooling all samples from one plate, a second PCR is performed to add two plate-specific barcodes (BC_plate_) as well as adapters required for Illumina sequencing (a1 and a2). Subsequently, all samples are pooled and sequenced via paired-end reading to cover all barcodes and mutation sites. **c**, Cell-specific hydroamination activity of 2,164 ArM variants from the initial library obtained by automated screening of 32 96-well plates. Only variants that were included for model training are displayed. Controls (empty vector and reference variant) are displayed with their standard deviation in red. **d**, Fraction of amino acids at the five randomized positions in Sav. Note that the amino acids of the reference variant (Sav 111T 112F 118N 119A 121Q, abbreviated Sav TFNAQ) are the most abundant, as the library was derived from this variant and contained at most four amino acid substitutions per variant.

Besides the activity assay, another critical barrier to obtaining sufficiently large sets of sequence-activity data can be the cost of sequencing. Obtaining the sequences of several thousand protein variants by Sanger sequencing typically costs more than USD 10,000, which is prohibitive for most academic labs. In principle, the cost per variant can be reduced significantly by relying on NGS, which quickly becomes more cost-efficient than Sanger sequencing as the library size increases. However, in NGS all variants are sequenced in bulk, which means a method to retroactively link each sequence to the corresponding activity measurement is required. Previously, the use of DNA barcodes has been suggested to enable NGS of protein variants distributed across 96-well plates^43–45^. Building on these strategies, we established a two-step PCR protocol for the barcoding of Sav variants that is compatible with the Illumina NGS platforms (Fig. 2b). In the first step, which is carried out in 96-well plates, the randomized region of the Sav gene is amplified using primers that append a well-specific barcode combination as well as constant regions to the ends of the PCR products. This is achieved using eight forward (representing the plate‘s rows) and twelve reverse primers (representing the columns). For simplicity, heat-treated samples of bacterial cultures serve as templates, avoiding the need for laborious and costly plasmid purification.

Subsequently, PCR products are pooled by plate, and each pool is gel-purified and used as a template for a second PCR. In this step, primers binding to the previously added terminal constant regions are used for amplification. These primers contain overhangs to append plate-specific barcodes as well as the adapters required for NGS. Through the combination of well-(1^st^ step) and plate-specific (2^nd^ step) barcodes, it is possible to sequence thousands of variants from multiple plates in a single, low-cost NGS run and to assign the obtained sequences to the corresponding activity value obtained in the functional assay. In our specific case, paired-end sequencing of 40 bp from one end and 110 bp from the other end of the final PCR product was sufficient to read all well- and plate-specific barcodes as well as the five mutation sites in the Sav gene at a high read coverage (average of >100-fold per variant) and low cost (see Discussion).

Relying on the combination of automated activity assay and NGS, we screened 32 96-well plates containing variants from the aforementioned library of Sav. As each plate contained six controls (empty vector and reference variant in triplicate), this amounts to a total of 2,880 variants. Excluding mutants that failed to grow, we obtained activity data on 2,790 variants. Most of these displayed an intermediate activity between the background level of cells lacking Sav (empty vector) and the reference variant Sav S112F K121Q (Fig. 2c). Notably, approximately 3 % of all mutants were more active than the reference. Using the NGS-based strategy, we retrieved the sequences for 2,663 out of 2,880 wells containing Sav mutants. After excluding variants with nonsense mutations and wells containing more than one variant, sequence-activity data for 2,164 clones were obtained, of which 2,035 were distinct variants. Notably, for variants appearing in multiple wells, the deviation between these replicate activity measurements was generally low, corroborating the high robustness of the assay (Supplementary Fig. 3). Importantly, the library displayed a high degree of sequence diversity, with every amino acid appearing in every position (Fig. 2d) and an average Hamming distance of 4.3 between the mutants. Note that the amino acids of the reference variant were the most abundant in each position, as we did not randomize all five positions simultaneously. Thus, the library exhibited a high degree of variability both in terms of activity distribution (including a low fraction of inactive variants) as well as sequence diversity. This indicated that the aforementioned design goals for the library were met, providing a promising data basis for modelling the sequence-activity landscape by machine learning.

As we had previously recorded sequence-activity data for 400 Sav double mutants (S112X K121X) that are part of the same sequence space^40^, we added these older data to the measurements obtained herein. As a result, a total of 2,992 data points covering 2,435 distinct ArM variants were available as initial training data for machine learning.

### Development of an initial machine learning model of ArM activity

To construct a model that can reliably predict the activity of untested ArM variants and guide further screening rounds, we relied on Gaussian process (GP) regression^46^. This machine learning technique can capture highly non-linear relationships and has the distinct advantage of being probabilistic, which means that it predicts a probability distribution rather than a point estimate, and thus provides an estimate for the confidence of each prediction. This feature can not only help users assess the uncertainty of individual predictions, but is ideally suited for active learning strategies. In this scenario, the model’s uncertainty estimates can be used to guide subsequent screening rounds towards uncertain regions of sequence space with the goal of improving the model (i.e., exploration), before suggesting highly active variants in later rounds (i.e., exploitation).

GPs are characterized by a mean and a covariance function, which is commonly referred to as kernel. In our case, as we operate on the space of protein sequences, the kernel measures the similarity between different ArM variants. Since the selection of a suitable kernel is of paramount importance for good performance and sample efficiency (i.e., predicting accurately with little data), we performed a benchmarking process and found that the non-linear Matérn kernel^46^ performed best in our case (see Methods).

Moreover, our model development pipeline included steps to account for experimental noise and to select suitable descriptors (Fig. 3a). Considering the inherent noise in biological experiments during modelling is crucial to ensure that decisions are not influenced by random fluctuations. To distinguish the genuine signal from these fluctuations, it is necessary to define a probabilistic model for data generation, known as the likelihood. This step involves specifying the likelihood and its parameters, which is essential for applying Bayes’ theorem to calculate the posterior distribution (see Methods). To elucidate the form of the likelihood, we relied on the variants appearing multiple times in the screening. This revealed that the deviation of these replicates from the per-variant mean closely follows a log-normal distribution, which can be viewed as a conservative estimate of the experimental noise in the data (Fig. 3b). Considering the log-transformed values, this implies a Gaussian likelihood. Next, we used the replicate measurements to determine a standard deviation, which is a key element in defining the data likelihood. We made the simplifying assumption that the variance of the measurement remains constant across the different ArM variants and repeated this analysis after each round of screening. As illustrated in Fig. 3c, under- or overestimating the experimental noise leads to a drastically reduced performance of the resulting model, likely due to overfitting to noise in the data. In contrast, the procedure applied here results in a robust performance in the face of noisy data.

**Fig. 3.**
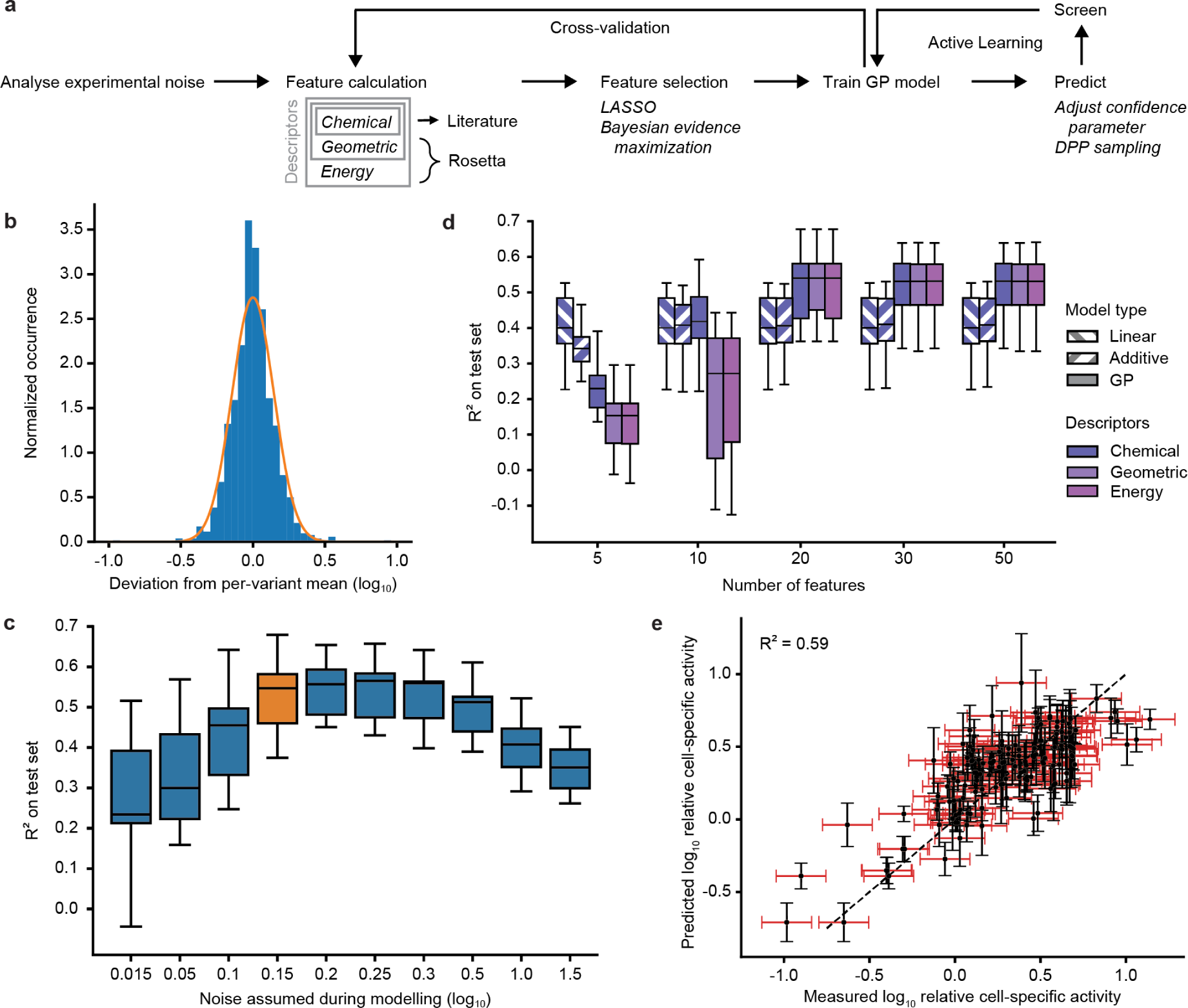
Development of the initial GP model. **a,** Overview of the machine learning pipeline. Initially, the standard deviation of the activity measurements was estimated to account for experimental noise. Subsequently, three feature sets were calculated and reduced sets were obtained by applying LASSO and Bayesian evidence maximization. The resulting descriptors were then used to train GP models. Model selection and model fitting were benchmarked using cross-validation. Ultimately, the GP model can be used to navigate the sequence space in active learning cycles. **b**, Histogram of the deviation between replicates in the initial library. The distribution of residuals can be conservatively approximated by a normal distribution with a specific variance (orange). **c**, Influence of the noise estimate on the predictive performance of the resulting GP model. The value chosen based on Fig. 3b is highlighted in orange. The models used here were based on chemical descriptors with 20 features (see Fig. 3d) and were evaluated using 15-fold cross-validation. The box plots display the 25th, 50th and 75th percentile with whiskers denoting the 1.5-fold interquartile range. **d**, Influence of feature number (x-axis), model type (fill pattern), and descriptors (colour) on the performance of machine learning models analysed by 15-fold cross-validation. The box plots display the 25th, 50th and 75th percentile with whiskers denoting the 1.5-fold interquartile range. **e**, Performance of the GP model using chemical descriptors and 20 features on an exemplary cross-validation split. The measurement uncertainty (one standard deviation) is displayed in red, while the uncertainty of the model is in black. The R^2^ value of this particular cross-validation split is displayed.

With regard to the descriptors that represent the ArM variants during training, we considered features that reflect chemical properties of amino acids^11^ as well as features that were extracted from Sav mutant structures predicted with the Rosetta software^47^. The latter included both geometric features (e.g. solvent accessible surface area, number of hydrogen bonds, partial charge, dihedral angles, etc.) and energy terms. Note that the geometric descriptors were compiled to be strict supersets of the chemical descriptors (i.e., they also included the chemical descriptors), and similarly the energy-based descriptors are strict supersets of the geometric descriptors. Given the large number of features (125 chemical, 682 geometric, and 161 energy features), we sought to select subsets that are parsimonious while still highly predictive to ensure data efficiency and eliminate redundancy. To this end, we relied on Bayesian evidence maximization (see Methods). Due to the non-linearity of the optimization challenge, we first reduced the feature sets using LASSO, which performed best in a benchmarking test (Supplementary Fig. 4). More precisely, we fitted a linear model and selected features with non-zero coefficients for automatic relevance detection using Bayesian evidence maximization with a Gaussian process. This allowed us to reduce the initial pool of features to 20-100 and speed up the evidence maximization step, which required multiple optimisation restarts to ensure that an adequate maximum was achieved.

Finally, we trained GP models using the different reduced feature sets on the available sequence-activity data and evaluated model performance using 15-fold cross-validation. For comparison, we included a linear and an additive, non-linear model based on chemical descriptors. The latter is restricted to treating potentially non-linear effects on the activity additively and is therefore not capable of modelling epistatic effects. Notably, the linear and additive models performed considerably worse than the GP models (Fig. 3d), confirming that advanced methods such as GP models are required to accurately capture the sequence-activity relationships in the data. Interestingly, the chemical, geometric, and energy-based descriptors displayed a comparable performance, and a set of 20 features proved to be sufficient in all cases. The most influential features based on automatic relevance detection are listed in Supplementary Table 1 (see Supplementary Fig. 5 for an analysis of their influence).

As computationally expensive structural calculations are required to generate the geometric and energy-based features and no clear benefit over models relying only on chemical descriptors was observable, we chose to continue with the subset of 20 chemical features as our primary encoding strategy for further modelling. The resulting model displayed a good predictive performance, with a median R^2^ of 0.54 based on 15-fold cross-validation (see Fig. 3e and Supplementary Fig. 6 for exemplary validation splits). While leaving room for improvement, this degree of correlation has previously been shown to be suitable for guiding directed evolution campaigns^11^. Moreover, the median Spearman correlation of 0.68 demonstrates that the relative ranking of variants was largely reproduced by the model (Supplementary Fig. 7), which is important for confident selection of high-activity variants.

### Model refinement by active learning

The aforementioned performance parameters indicate that the initial GP model can predict ArM activity with reasonable accuracy. However, due to the vast sequence space, the random sampling from this space during the generation of training data, as well as inevitable biases in experimental library construction, it is likely that this initial model will not generalize well across the entire sequence-activity landscape. Consequently, it may be “blind” for certain underexplored regions containing highly active ArMs. Therefore, we performed a second, exploratory screening round with the goal of improving the model’s accuracy and ability to generalize across the entire sequence space. To this end, we designed a new library consisting of 720 variants that were primarily selected to be “informative”. Specifically, we utilised the uncertainty estimates of the GP model and selected the variants with the highest uncertainty in the predicted activity among all 3.2 million mutants^48,49^.

We generated these variants based on a pool of oligonucleotides obtained through commercial synthesis on arrays, a method that allows for the cost-efficient construction of large and targeted libraries^50^ and is therefore highly useful for active learning with large batch sizes. After cloning the oligonucleotides into the Sav expression plasmid, we screened the resulting exploration library relying on the automated pipeline in combination with NGS as described above. This exploratory round yielded sequence-activity data on 465 additional variants. It should be noted that this library also contained chimeric variants with amino acid combinations that were not planned in the computational design, likely due to PCR-mediated recombination between variants^51,52^. While unintended, these additional variants can also be used to augment the machine learning model and were therefore included for training. If desired, chimera formation can be minimized by optimizing the PCR conditions^51,52^.

The exploration library displayed a similar activity distribution as the initial training data (Fig. 4a), which is in line with the focus on informative instead of active variants. Importantly, these new data led to a decrease in the standard deviation of the predictions, most prominently for variants that had previously exhibited a high uncertainty (Fig. 4b). While this observation alone is not a proof of increased accuracy, it hints towards an improved representation of previously underexplored regions of the sequence space, which we examined in more detail in subsequent analyses (see below).

**Fig. 4.**
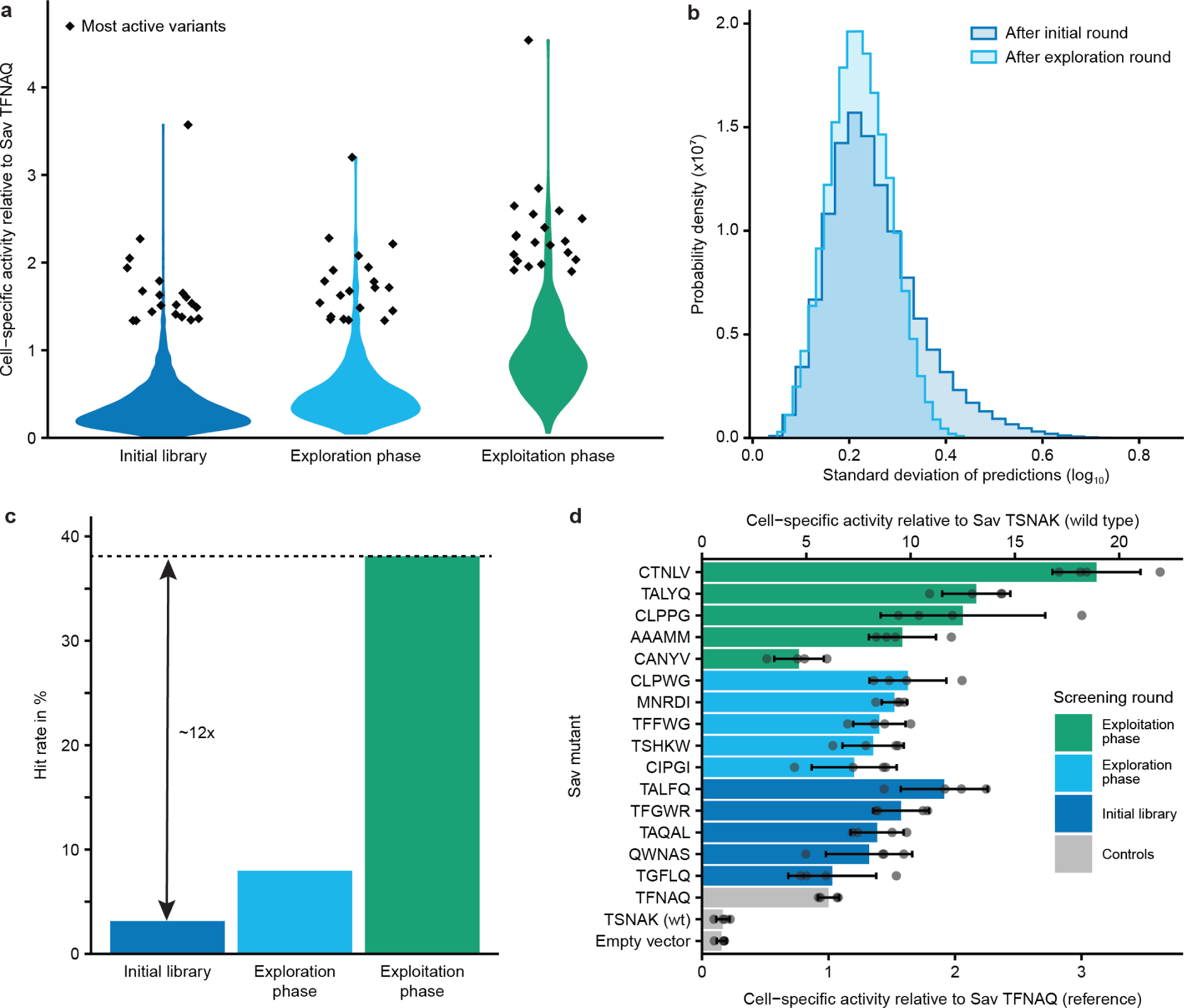
ArM engineering by means of active learning. **a**, Activity distributions in the three screening rounds displayed as violin plots. The 20 most active variants in each round are depicted as diamonds. Activity is displayed relative to the reference variant (Sav TFNAQ). **b**, Normalized histograms of the standard deviations of predictions across all 3.2 million variants after the first and second round of screening. **c**, Hit rate in the three screening rounds. Here, any variant with a higher cell-specific activity than the reference variant is considered a hit. The hit rate represents the fraction of hits amongst all variants screened in the respective round. Note that the hit rate in the initial library was calculated based on the triple and quadruple mutants, excluding the double mutants that had been tested previously^40^. In the third round, chimeric variants that were not part of the computationally designed library were excluded to provide a better analysis of the models’ performance. **d**, The five most active variants from each screening round were tested again in four replicates. The five-letter codes denote the amino acids in positions 111, 112, 118, 119, and 121 for the respective variants.

### Active learning increases the efficiency of directed evolution

Following model refinement in the exploration round, we set out to test whether our model-guided approach can indeed aid in the discovery of active ArMs. With this goal in mind, we designed a third library of 720 variants predicted to be of high activity. Additionally, we employed an *in silico* diversification step to avoid choosing only variants with highly similar sequences. This provides a safeguard against inaccuracies in the top predictions and increases the likelihood of obtaining variants with diverse properties besides activity (e.g. thermostability, solubility, or activity under alternative conditions). To this end, we used a notion of diversity known as determinantal point processes (DPPs)^48,49^, which use the GP kernel to determine which variants are similar to each other (see Methods and Supplementary Fig. 8a). In short, this approach treats the descriptors of the Sav variants as vectors in Euclidian space and attempts to select a set of vectors that are as orthogonal to each other as possible. We applied this process to a set of variants with the highest predicted activity to obtain a subset of active and yet sequence-diverse variants. This led to a more diverse set of variants compared to a simple greedy selection of the variants with the highest predicted activity as assessed by three different metrics of diversity (Supplementary Fig. 8b).

As described for the exploration round, we obtained the designed library based on an oligonucleotide pool and acquired experimental data for 349 distinct variants. Gratifyingly, this third library displayed a clear shift towards higher activities compared to the first two rounds, both in terms of the average as well as the top activities (Fig. 4a). We further analysed the hit rate in the screening rounds, which we define here as the fraction of ArM variants with higher activity than the reference variant, which is the most active variant identified in a previous study^40^. While only 3 % of the initial library were hits, this rate reached 38 % in the exploitation phase, amounting to an approximately 12-fold increase (Fig. 4c). This demonstrates that the model acquired a meaningful representation of the activity landscape and can reliably predict active ArMs.

To confirm the results from the different screening rounds, which were performed in single measurements, we tested the most promising variants from all three rounds again in four replicates (Fig. 4d). This revealed that Sav 111C 112T 118N 119L 121V (abbreviated Sav CTNLV) was the most active variant, reaching an 18-fold higher cell-specific hydroamination activity than the wild type (Sav TSNAK) and a three-fold higher cell-specific activity than the reference variant (Sav TFNAQ). In addition, we purified the most active variants from our whole-cell screening to test whether they also display an increased total turnover number *in vitro*, which was the case for five of the seven variants tested (Supplementary Fig. 9). As observed before^40^, the ranking of the variants changed *in vitro*, which can be expected due to the different reaction environments and varying expression levels in the periplasmic screening.

Notably, the Sav CTNLV mutant does not retain the S112F K121Q mutations that were found to be optimal in the previous double mutant screening^40^. Likewise, all other variants evaluated in the validation experiment (Fig. 4d) retain neither or only one of these two mutations. This highlights the importance of epistatic effects, which can only be adequately considered through combinatorial library designs and non-additive models. Strikingly, several highly active variants contain a cysteine at position 111, which seems counter-intuitive as cysteine has been repeatedly shown to have a pronounced inhibitory effect on gold-catalysed hydroamination^53^. However, residue 111 is pointed away from the metal, presumably preventing the thiol from interfering with catalysis. Notably, the beneficial impact of this mutation was not obvious from the initial data set, but became increasingly apparent in subsequent rounds. This indicates that active learning can traverse the mutational space more broadly than alternative methods and enable the identification of counter-intuitive effects on activity.

To further corroborate this hypothesis, we performed more detailed analyses to investigate whether the active learning strategy with a model-guided exploration round indeed led to a better representation of the available sequence space. We visualised the sequence space (using t-SNE^54^ on the kernel matrix, see Methods) to analyse how the tested variants are distributed across this space (Fig. 5a, b). While care must be taken when interpreting such low-dimensional projections, this analysis indicates that the initial library did indeed not cover the sequence space uniformly. The subsequent exploration round filled in several of the “gaps” in accordance with the design goal of this phase. The exploitation phase focused on a number of regions of high activity, indicating that the selection criteria of high activity and sequence diversity were met. The emergence of multiple clusters of active variants is compatible with the notion of a “rugged” activity landscape with many local optima. Such landscapes can be challenging to navigate using classical methodologies, which frequently follow a single “uphill” trajectory. In contrast, the GP model developed here acquires a holistic understanding of the entire space of 3.2 million ArM variants and allows us to sample various potential optima, increasing the chances of finding suitable variants.

**Fig. 5.**
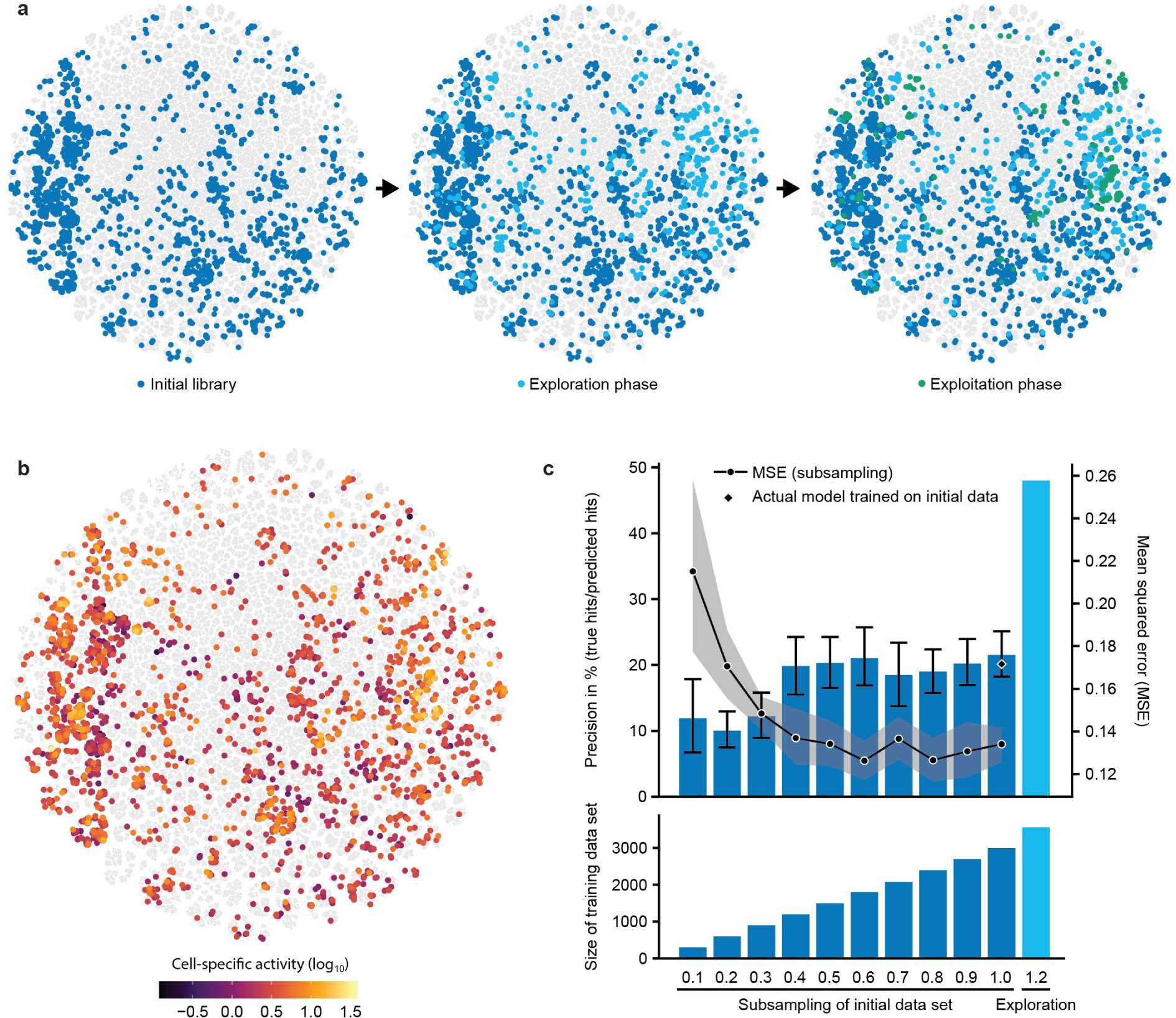
Enhanced sequence-activity mapping through active learning. **a,** t-SNE visualisation of the sequence space. ArM variants that were tested in the three screening rounds are highlighted in different colours. To generate this visualisation, all 3.2 million mutants were considered, and a uniform subsample of untested variants was plotted in grey. The similarity metric used was derived from the GP model (see Methods for details). **b**, t-SNE visualisation of the sequence space with colour encoding the activity of experimentally tested variants. The clustering is identical to that in Fig. 5a. **c**, Precision in identifying hits and mean squared error (MSE) of predictions as a function of the size of the training data set. The dark-blue bars in the upper graph indicate the average precision of models that were trained on different fractions of the initial data set (screening round 1). The diamond at 1.0 represents the precision of the model used to inform experiments. The light-blue bar on the right represents the model refined by model-guided exploration (screening round 2). Note that the precision is not identical to the experimentally determined hit rate (see Methods). The lower graph depicts the size of the data sets used to train the respective models.

Lastly, we sought to quantify the effect of the applied sampling strategy in relation to the size of the training data set. A crucial question in this regard is whether the active learning strategy suggested here provides a significant benefit over a comparable increase in the size of the training data set by random sampling of variants. To investigate this, we trained models on different fractions of the initial data set using the same model development pipeline as before. As a proxy for an experimentally determined hit rate, we analysed the models’ precision in identifying hits among the variants tested in the exploitation phase (i.e., the percentage of true hits among variants predicted to be hits). As illustrated in Fig. 5c, this analysis indicates that acquiring training data by random sampling is accompanied by strong diminishing returns: Approximately 40 % of the initial data set size (equivalent to ∼1200 data points) is sufficient to achieve a similar performance (in terms of precision and mean squared error (MSE)) as a model trained on the entire initial data set (∼3000 data points). This suggests that additional random screening rounds of similar size would not have led to noteworthy improvements of the model. In contrast, the model-guided exploration round, which consisted of only 564 additional data points (an increase of less than 20 % in data volume), improved the precision in identifying hits from ∼20 % to 48 %. This increase is significantly beyond any improvement that can be anticipated due to the mere increase in data volume, emphasizing the fact that this round was substantially more informative than random sampling. This confirms the validity of the suggested active learning and model-guided exploration strategies, pointing to a high potential for enhancing MLDE campaigns while at the same time minimizing the experimental effort.

## Discussion

MLDE is a highly promising strategy for engineering enzymes and other proteins. However, the success and efficiency of such engineering campaigns hinges on the ability to generate sufficiently large and informative data sets, the use of smart sampling strategies, and the choice of suitable machine learning techniques that optimally leverage the resulting data.

Many studies on MLDE have relied on small data sets^4–10^ and a single training phase^4,5,10,55,56^, which may be attributed to experimental limitations. This bears the risk that the resulting models do not accurately represent the sequence space, and thus are likely to leave significant potential hidden within this space untapped. Here, we applied lab automation and NGS to acquire large data sets in a simple and cost-efficient manner, and directed our sampling to the most informative data by means of advanced active learning techniques.

Lab automation greatly increases the throughput of screenings and is, at the same time, highly adaptable to various reactions and target proteins. In this study, we performed some experimental steps manually, but a fully automated workflow could also be implemented. Similarly, the computational pipeline is largely automated, and thus it is conceivable to conduct protein engineering with minimal human intervention. Importantly, recent developments such as academic biofoundries and cloud labs are making such approaches more widely accessible^57,58^.

The NGS strategy employed here enables the sequencing of thousands of protein variants for the cost of a small Illumina run and PCR reagents. The former is available for a few hundred dollars (e.g. MiSeq Nano, yielding approx. 1 million reads) and will likely continue to get cheaper. If combined with other samples and run on an instrument with a large capacity, the prorated costs may even be in the range of a few dollars. Regarding the PCR reagents, primer synthesis costs are low as only 20 primers are required to address all 96 positions in a well plate. Similarly, the use of two plate barcodes means that 12 primers for the second PCR are sufficient to distinguish 36 well plates. Overall, this means that sequencing is possible at a cost of less than one cent per variant.

Combined, automation and NGS are ideally suited to generate large data sets for MLDE. At the same time, it is also crucial to design information-dense libraries to maximize the efficiency of experimental screening rounds. In the initial round, we achieved this by optimizing the mutational load in the library, which is a straightforward and broadly applicable strategy. Alternatively, approaches such as zero-shot methods relying on ΔΔG calculations^13^ can be applied as well. In subsequent rounds, library design can be guided by the machine learning model. While it may seem attractive to apply an exploitation-focused strategy to quickly identify active variants, we hypothesized that a model-guided exploration round could substantially improve the predictive performance and thus increase the chances of identifying suitable variants in large and rugged sequence spaces in a subsequent round. Indeed, we observed that the exploration round improved the model’s ability to identify active variants far beyond what would be expected due to the increase in data volume alone. This demonstrates that active learning is a highly effective and efficient strategy for developing accurate models of sequence-activity landscapes. Moreover, the separation into exploration and exploitation phases provides a transparent and practical solution to the exploration-exploitation dilemma, as it allows for a clear and plannable resource allocation. In addition, our study introduces DPP sampling as a strategy for diversifying the selection of active variants, which increases the robustness of MLDE to possible model inaccuracies and may be beneficial with regard to secondary properties beyond activity.

In terms of the machine learning approach, this study corroborates that Gaussian process regression is an attractive choice for MLDE, particularly when strong epistatic effects are present in the sequence-activity landscape. Moreover, it is well-suited for active learning strategies, as the uncertainty quantification is computationally simple, which constitutes an advantage over alternative methods such as deep learning. Our results demonstrate that simple and computationally efficient descriptors are sufficient for non-trivial improvements to engineering campaigns, which is in line with other literature on the subject^59,60^. Nonetheless, it might be possible to further boost the predictive performance, for example by employing improved structure prediction algorithms or descriptors from modern protein language models^61,62^. Lastly, our results highlight that accurately accounting for experimental noise is crucial during model development, an aspect that has frequently been neglected^63^.

The application of these strategies to the engineering of ArMs for gold-catalysed hydroamination led to the identification of a variant with 18-fold higher cell-specific activity than the wild type. Compared to our previous screening of double mutants^40^, extending the search space to five positions led to a three-fold improvement. Further rounds of active learning could potentially lead to the discovery of even more active variants. Moreover, the methods developed here could be used to target additional positions. However, it should be noted that this ArM is likely a challenging engineering target due to the relatively exposed location of the cofactor in Sav. Therefore, applying this engineering strategy to alternative scaffolds with a more shielded active site might enable larger improvements^64^. Currently, artificial (metallo)enzymes are typically limited by their rather modest activity. Thus, the field could profit greatly from advanced machine learning-guided engineering strategies, as demonstrated here. Similarly, the active learning approach described here could be applied to tailor natural enzymes for industrial applications, or to engineer other proteins such as antibodies, biosensors, or transporters.

## Materials and Methods

### Chemicals and reagents

**(Biot-NHC)Au1** was synthesized as previously described^40^. All other chemicals were obtained from Sigma-Aldrich. Primers were synthesized by Sigma-Aldrich, and enzymes for molecular cloning were obtained from New England Biolabs.

### Plasmids

All plasmids were based on a previously described expression plasmid that contains a T7-tagged Sav gene with an N-terminal OmpA signal peptide for export to the periplasm under control of the T7 promoter in a pET30b vector^33^. This plasmid is available from Addgene (#138589). A version of this plasmid encoding the Sav S112F K121Q mutant was used as the starting point for library generation.

### Cloning of Sav libraries

#### Site-saturation mutagenesis at 20 positions

To individually randomize 20 positions in Sav, the plasmid encoding Sav S112F K121Q was amplified in two parts in order to create two overlapping fragments for each position, with mutations being introduced by an NDT codon in one of the primer overhangs. The PCRs were conducted using the primer pairs SSM_X_NDT_fwd and kanR_rev, and kanR_fwd and SSM_X_rev (X denotes the position to be randomized, see Supplementary Table 2). PCRs were carried out using Q5 High-Fidelity DNA Polymerase (New England Biolabs). Following DpnI digest and PCR purification, the corresponding fragments were assembled by Gibson assembly and transformed into *E. coli* BL21-Gold(DE3). Three clones per position were sequenced by Sanger sequencing to verify correct assembly and diversity at the desired position.

#### Double, triple, quadruple, and quintuple mutant libraries

To generate sets of double, triple, quadruple, and quintuple mutants, the plasmid encoding Sav S112F K121Q was amplified in two parts. One part included the Sav positions 111 and 112, and the other part included positions 118, 119, and 121. To generate fragments with variable but defined numbers of mutations, the primers from Supplementary Table 3 were used in several PCR reactions according to Supplementary Table 4. Following DpnI digest and PCR purification, the fragments were assembled in several Gibson assembly reactions as summarized in Supplementary Table 5. The reactions were then transformed separately into chemocompetent *E. coli* Top10. Plasmids were isolated from the transformants and transformed into the expression strain BL21-Gold(DE3). When picking colonies for screening, the theoretical diversity of the individual sub-libraries (Supplementary Table 5) was taken into account in order to obtain balanced sets of double, triple, quadruple and quintuple mutants.

#### Active learning libraries

To create libraries of specific Sav variants that were suggested by the machine learning models, oligo pools were ordered from Twist Bioscience. These oligos were used as primers that bind immediately downstream of position 121 in Sav. The 5’-overhang contained the five mutation sites with the desired changes as well as a constant region for Gibson assembly (see Supplementary Table 6). For the first library of ML-designed variants, insert and backbone were generated according to Supplementary Table 7. For the second library, the PCRs were run according to Supplementary Table 8. Following DpnI digest and PCR purification, the fragments were assembled by Gibson assembly and transformed into chemocompetent *E. coli* Top10. Plasmids were isolated from the transformants and transformed into the expression strain BL21-Gold(DE3).

### Sav expression in 96-well plates

96-deep well plates were filled with 500 µL of LB (+ 50 mg L^−1^ kanamycin) per well. Cultures were inoculated from glycerol stocks and grown overnight at 37 °C and 300 revolutions per minute (rpm) in a Kuhner LT-X shaker (50-mm shaking diameter). 20 µL per culture was used to inoculate expression cultures in 1 mL of LB with kanamycin. These cultures were grown at 37 °C and 300 rpm for 1.5 h. At this point, the plates were placed at room temperature for 20 min, and subsequently, Sav expression was induced by addition of isopropyl-β-D-thiogalactopyranoside (IPTG, final concentration 50 µM). Expression was carried out at 20 °C and 300 rpm for an additional 16 h.

### Whole-cell screening

Following the expression of Sav mutants in deep-well plates, the OD_600_ of the cultures was determined in a plate reader using 50 µL of samples diluted with an equal volume of PBS. Afterwards, the plates were centrifuged (3,220 rcf, 15 °C, 10 min), the supernatant was discarded and the pellets were resuspended in 400 µL of incubation buffer (10 µM **(Biot-NHC)Au1** in 50 mM MES, 0.9 % NaCl, 10 mM diamide, pH 6.1). Cells were incubated with the cofactor for 1 h at 15 °C and 300 rpm. Afterwards, plates were centrifuged (2,000 rcf, 15 °C, 10 min), the supernatant was removed and the pellets were resuspended in 500 µL of washing buffer (50 mM MES, 0.9 % NaCl, 10 mM diamide, pH 6.1). Following another centrifugation step, cell pellets were resuspended in 200 µL of reaction buffer (5 mM 2-ethynylaniline in 50 mM MES, 0.9 % NaCl, 10 mM diamide, pH 6.1). Reactions were performed at 37 °C and 300 rpm for 20 h before determining the product concentration. To account for differences in cell density and plate-to-plate variations, the product concentrations were divided by the OD_600_ of the culture and normalized to the mean of the cell-specific product concentrations measured for the Sav S112F K121Q controls in the respective plate.

### Kovac’s assay

Indole was quantified using the photometric Kovac’s assay (adapted from Piñero-Fernandez et al.^65^). For measurements in culture supernatant, plates were centrifuged (3,220 rcf, 20 °C, 10 min) and 110 µL supernatant was mixed with 165 µL of Kovac’s reagent (50 g L^-1^ 4-(dimethylamino)benzaldehyde, 710 g L^-1^ isoamyl alcohol, 240 g L^-1^ hydrochloric acid) in a separate plate. After 5 min of incubation, these plates were centrifuged (3,220 rcf, 20 °C, 10 min). Subsequently, 75 µL of the upper phase was transferred to a new transparent plate and the absorbance at 540 nm was measured in a plate reader (Tecan Infinite M1000 PRO).

### Lab automation

Colony picking, reaction setup and product quantification were implemented using an automation platform featuring two Tecan EVO 200 (Tecan Group AG) robotic platforms coupled to each other. Both platforms were controlled using the EVOware standard software (Tecan Group AG). Colony picking was performed using the integrated Pickolo system (SciRobotics). For shaking, incubation, and resuspension of cultures, the platform was equipped with a Kuhner ES-X shaking platform (Adolf Kühner AG) running at 300 rpm at 50-mm shaking radius. The shaking platform was surrounded by a custom-made box made of aluminum plastic composite panels (Tecan Group AG). The temperature inside the box was maintained at 15 °C using an “Icecube” (Life imaging services) heater/cooler device. Centrifugation of the samples was performed using the integrated Rotanta 46 RSC Robotic centrifuge (Hettich AG). All buffer exchanges during sample preparation were performed using the integrated liquid-displacement pipetting system equipped with eight 2500 µL dilutors and fixed stainless steel needles. Absorbance measurements were performed using a Tecan Infinite M200 PRO plate reader. The automation method files are available upon request.

### Barcoding of mutants

Following colony picking, cultures were grown overnight at 37 °C and 200 rpm in 96-deep well plates. On the following day, 150 μL per culture was transferred to a 96-well PCR plate. The plates were sealed and placed in a thermal cycler for 5 min at 95 °C to lyse the bacteria. Subsequently, the plates were centrifuged (3200 rcf, 5 min) and 0.5 μL of the supernatant was used as template for the first PCR. This PCR step was done in 96-well plates, with each well containing a distinct combination of barcoded primers (see Supplementary Table 9). 30 cycles were performed with 30 s denaturation at 98 °C, 20 s annealing at 71 °C and 30 s elongation by Pfu DNA polymerase at 72 °C. The products from each plate were pooled, run on a 2.5 % (w/v) agarose gel at 100 V for 2 h and purified using a gel extraction kit (Sigma-Aldrich). The products were then used as templates for a second PCR with distinct combinations of barcoded primers (Supplementary Table 10) to generate a plate-specific labelling. The primer overhangs also contained the adapters required for Illumina sequencing. 30 additional cycles were performed, consisting of 30 s denaturation at 98 °C, 20 s annealing at 63 °C and 30 s elongation by Q5 High-Fidelity DNA Polymerase (New England Biolabs) at 72 °C. Ultimately, all products were pooled, run on a 2.5 % (w/v) agarose gel at 100 V for 2 h, and purified using a gel extraction kit.

### Illumina sequencing

NGS was performed by the Genomics Facility Basel using an Illumina MiSeq platform and a Reagent Kit v2 Nano (150 cycles, PE 110/40) using ∼20 % genomic PhiX library as spike-in to increase sequence diversity.

### NGS data analysis

NGS data were analyzed using a custom R script. Forward and reverse reads retrieved from fastq files were paired and target fragments were selected based on several constant regions (GTCACACGTAGCATGTGG, GAGACCTTGTGTCGATGG, GGCCTCGGTGGTGCC, no mismatches). Mutation sites as well as barcodes were extracted based on their distance to these regions. All reads with a Q-score < 30 at the mutation sites were discarded, as well as those for which a barcode did not match any of the expected sequences. The codons at the mutation sites were translated to amino acids in order to identify the Sav variants and the barcodes were used to identify the plate and well for each read. For each plate, the entries were then grouped by variant and only the combinations of variant and well with the highest number of reads was kept. This eliminates combinations of variants and barcodes that result from chimera formation during the second PCR step. Subsequently, variants that accounted for less than 80 % of reads for a given barcode combination were discarded in order to eliminate cases where more than one variant had been present in a well.

### Sav expression for purification

A single colony of *E. coli* BL21-Gold(DE3) harbouring a plasmid for periplasmic expression of the desired Sav variant was used to inoculate a starter culture (4 mL of LB with 50 mg L^-1^ kanamycin), which was grown overnight at 37 °C and 200 rpm. On the following day, 100 mL of LB with kanamycin in a 500 mL flask was inoculated to an OD_600_ of 0.01. The culture was grown at 37 °C and 200 rpm until it reached an OD_600_ of 0.5. At this point, the flask was placed at room temperature for 20 min and 50 µM IPTG (final concentration) was added to induce Sav expression. Expression was performed at 20 °C and 200 rpm overnight, and cells were harvested by centrifugation (3,220 rcf, 4 °C, 15 min). Pellets were stored at −20 °C until purification.

### Sav purification

Cell pellets were resuspended in 10 mL of lysis buffer (50 mM tris, 150 mM NaCl, 1 g L^−1^ lysozyme, pH 7.4). After 30 min of incubation at room temperature, cell suspensions were subjected to three freeze-thaw cycles. Subsequently, nucleic acids were digested by addition of 10 µL of DNaseI (2000 units/mL, New England Biolabs) and CaCl_2_ to a final concentration of 10 mM, followed by incubation at 37 °C for 45 min. After centrifugation, the supernatant was transferred to a new tube and mixed with 40 mL of binding buffer (50 mM ammonium bicarbonate, 500 mM NaCl, pH 11). Pierce iminobiotin agarose (Thermo Fisher Scientific) was equilibrated in falcon tubes and used to pack a PD-10 column up to a bed height of approximately 1 cm. The lysate was loaded onto the column relying on gravity flow. Subsequently, the column was washed twice with 10 mL binding buffer. Ultimately, Sav was eluted using 10 mL of elution buffer (50 mM ammonium acetate, 500 mM NaCl, pH 4). Amicon Ultra filters (10 kDa molecular weight cut-off) were then used to concentrate the samples and exchange the buffer against the reaction buffer (50 mM MES, 0.9 % NaCl, pH 6.1).

### Quantification of biotin-binding sites

The concentration of Sav biotin-binding sites was determined using a modified version of the assay described by Kada et al.^66^, which relies on the quenching of the fluorescence of a biotinylated fluorophore upon binding to Sav. Specifically, 190 µL of the binding site buffer (1 µM biotin-4-fluorescein, 0.1 g L^−1^ bovine serum albumin in PBS) was mixed with 10 µL of purified Sav. After incubation at room temperature for 90 min, the fluorescence intensity was measured (excitation at 485 nm, emission at 525 nm), and a calibration curve produced with lyophilized Sav was used to calculate the concentration of Sav biotin-binding sites.

### *In vitro* catalysis

*In vitro* reactions were performed with 2.5 µM purified Sav (tetrameric; corresponding to 10 µM biotin-binding sites), 5 µM **(Biot-NHC)Au1** and 5 mM 2-ethynylaniline in MES buffer (50 mM MES, 0.9 % NaCl, pH 6.1). The reactions were performed in a volume of 200 µL in glass vials and were incubated at 37 °C and 200 rpm for 20 h. Subsequently, the indole concentration was determined using the Kovac’s assay.

### Machine learning

All machine learning methods were implemented in Python using scikit-learn^67^, Pytorch^68^, Biotite^69^, pyRosetta^70^ and SciPy^71^.

#### Calculation of descriptors

In this work, we encoded the Sav mutants by three different classes of descriptors: chemical descriptors, geometric descriptors, and energy-based descriptors. To obtain the chemical descriptors, we utilized amino-acid descriptors from four different sources: Z-scores^16^, VSHE^17^, Barley score^18^, and PCscores^55^. All of these are based on physical amino-acid properties (see Supplementary Table 11) and principal component analysis (PCA) was used to construct a reduced representation. Here, we concatenated these features, resulting in 25 values per amino-acid position. As we considered quintuple mutants, each Sav variant is thus described by 125 features.

The geometric and energy-based features were created using the Rosetta software. First, we calculated the approximate dimeric structure of each mutant with a fixed seed using the *mutate* function with the default distance for post-mutational changes. The mutations were performed in the order of the five sites in the primary protein sequence (111, 112, 118, 119, 121). We calculated all 3.2 million approximate Sav dimer structures. Next, we used the package Biotite to calculate charge, distance to the centre of mass, and radii of each amino-acid residue. Additionally, we calculated the solvent accessible surface area of each residue, the number of hydrogen bonds per residue, and the dihedral angles. A summary of the features can be found in Supplementary Table 12. We discarded variables that did not vary across the 3.2 million structures, leaving us with 682 features. The energy-based features were calculated in the same manner as the geometric features using the approximate structure of the variant and correspond to the ref2015 set of 31 features per mutant from the Rosetta suite (see Supplementary Table 13). A common pre-processing step applied to all features involved subtracting the mean of each descriptor across the 3.2 million variants and scaling by the absolute value of the maximum value of that descriptor. This process ensured that the descriptors fell within the range [-1,1] and that their average value was zero.

#### Likelihood elucidation

The first step of any data analysis is to understand its randomness and generation process. In our case, the likelihood specified the experimental error introduced by biological variability, the measurement procedure, etc. In other words, we assumed that our measurements were corrupted by additive noise under log transformation. To justify this hypothesis, we analysed the distribution of the differences between replicates from their mean value. As a normal distribution appeared to be a good and conservative approximation for these data, we used a Gaussian likelihood with a standard deviation determined from the aforementioned distribution. In the first round, this value was determined to be 0.15, rounded to two decimal points in the log-transformed cell-specific activity. We repeated the same procedure for the subsequent screening rounds to account for variability between experiments. The standard deviations determined for the second and third round were 0.20 and 0.12, respectively.

#### Model section

For further analysis and Gaussian process fitting, we did not use the full set of features due to the complexity of the initial fitting procedure, which involves optimizing the marginal likelihood^72^. To simplify this process, we preprocessed the initial set of descriptors using one of three straightforward machine learning models: LASSO, elastic net, and random forests. We evaluated the effectiveness of this procedure through cross-validation on the entire feature space. In all cases, we utilized the scikit-learn implementation of these methods. Both the LASSO and elastic net methods employed an adaptive selection of the regularization parameter, which involved an additional layer of cross-validation within the training split. For random forests, we used a configuration of 500 trees with a maximum depth of 15 and a minimum split size of 5. After training, we selected *k* descriptors with either the largest coefficients or the highest feature importance for further analysis. We varied *k* across 20, 40, 60, 80, and 100. This range was chosen as the maximum set of descriptors that we believed would allow the Gaussian process library to reliably optimize the marginal likelihood.

#### Gaussian process

The functional relationship between the Sav sequence and ArM activity was modelled using Gaussian processes (GPs). This Bayesian method is versatile in capturing a wide range of structures, and is defined by its mean and covariance function, also known as the kernel. In our case, we found that kernels of the following form performed best among selected statistical models with calibrated uncertainty:

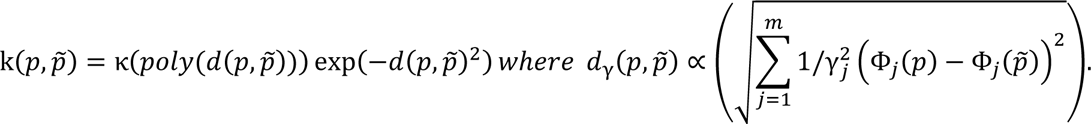

This kernel is known as Matérn kernel with regularity parameter η=5/2 and is commonly used to model twice differentiable smooth response surfaces^46^. The letters p and p’ denote different protein variants of which we want to calculate similarity. The function Φ corresponds to the feature representation of the protein p. In this work, this is a function that maps the protein sequence or structure to a fixed length vector. The parameters γ_*i*_ are usually referred to as length scales and are used for automatic relevance detection^73^. They guide the importance of a certain variable, i.e., if γ is very large, this part of the descriptor vector Φ has less impact if changed than a coordinate Φ_*j*_ with larger γ_*j*_. The length scales can be selected based on Bayesian evidence maximization, which is a well-tested methodology to select length scales that most likely explain the activity data^72^. The parameter κ was selected using the expected maximal achievable improvement of the protein, in this case κ = 3, meaning that the maximum achievable improvement is 1000-fold over the wild-type variant (due to modelling log_10_).

#### Bayesian evidence maximization

Hyperparameters, specifically the length scales of the Matérn kernel, were optimized for each of the chosen features using the maximization of evidence, a common Bayesian approach^46^. As before, we denote length scales *γ*. By maximization of evidence, we mean

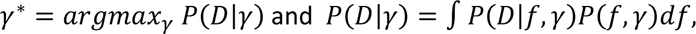

where *P*(*f*|*γ*) is the Gaussian process prior parametrized by length scales, and *P*(*D*|*f*,*γ*) is the Gaussian likelihood as specified in the prior section on likelihood elucidation. The integration in the prior formula represents marginalization of the Gaussian process, and strictly speaking integration requires certain mathematical regularity conditions, which we omit here. Upon finding the right length scales from the initial data, these were fixed, and the posterior *P*(*f*|*γ*,*D*) was calculated after each experimentation round without changing them. To implement the Bayesian posterior calculation, we used a custom implementation in Python.

### Active Learning

To employ active learning, we used a technique similar to the upper confidence bound method as described by Srinivas et al.^27^, or greedy information maximization. In the exploration round, we generated predictions using the GP model based on chemical descriptors with 20 features. To select informative variants, the confidence parameter was set to infinity. In addition, we allocated a smaller part of the experimental budget to variants predicted to be active to validate the model. The latter budget was split equally into three categories: A conservative set representing the Sav mutants for which the mean prediction minus two standard deviations was highest, as well as balanced and optimistic predictions chosen based on the mean and the mean plus two standard deviations as ranking mechanisms, respectively. See Supplementary Table 14 for an overview of the budget allocation in the exploration round. We obtained additional data points through a small random mutagenesis as well as chimeric variants, which were not part of the designed library.

In the exploitation round, we aimed to select active and diverse ArM variants. To this end, we trained three GP models on the new data set (including the exploration round). The three models employed different descriptors (chemical descriptors with 20 features, geometric and energy –based descriptors with 50 features) to possibly obtain more diverse predictions. We split the experimental budget equally among the three models. Further, we split the experimental budget per model into conservative and balanced predictions (see above). The experimental budget allocation can be found in Supplementary Table 15. The confidence parameter was set to 2 for the exploitation round. Additionally, a diversifying principle based on determinantal point processes^48^, a mathematical model of diversity, was employed to choose a diverse subset of variants, following the principles described by Nava et al.^49^ (see below). Upon retrieval of the above budget, we performed a validation step. As part of it, we augmented the chemical descriptor model with the new data and proposed 30 additional Sav variants to test for potential improvements. These were selected to be conservative or balanced (10 variants each), and 10 variants were selected to be the best predicted according to the balanced prediction metric.

### DPP sampling

When selecting Sav variants for experimental testing, it is advisable that these are diverse, especially in the context of the exploitation round. For example, if we were to identify the best *x* candidates using the machine learning pipeline, it is very likely that all these top *x* candidates are highly similar to each other for small *x*. If the model happens to be incorrect with regard to the top predictions, this will lead to failure to identify any active mutants. A more principled approach is to pick a diverse subset. Namely, select a set of promising mutants, and then further select a subset of these which is diverse. This ensures robustness to potential misspecification errors. The model of diversity we employed here is the inverse of the similarity model we used to train the GP regressor, namely the kernel. We measured the diversity of the selected subset by the determinant of the kernel matrix. This is a common approach in the machine learning literature^48^, as it has an intuitive interpretation where the determinant between two vectors is proportional to the volume that the two vectors span (see Supplementary Fig. 7a). The more orthogonal (dissimilar) these two vectors are, the larger the volume. A natural extension to non-parametric models such as GP models is to use the kernel matrix instead of the inner product between vectors. Finding a subset of maximum determinant is an NP-hard problem^74^. Hence, often a probabilistic method is employed to find the subsets^49,75^.

Suppose that the probability of sampling a set is proportional to the value of the determinant for this set. This probabilistic object is known as determinantal point process (DPP)^48^ and can be sampled very efficiently. In order to diversify our top-*x* batches, we select a top *y* number of candidates, where *y* is bigger than *x*, from which we choose a diverse set of size *x* using DPP sampling. The value of *y* = 500 was chosen arbitrarily for our experiments. The value of *x* depends on the available experimental budget in each round. The explorative round does not require diversification as the goal to select informative Sav variants already leads to diversity. In fact, it is related to the greedy search for a set with the largest determinant^75^.

In order to compare the diversity of the measurements, we use the isometry score, which is a ratio determinant and trace of a kernel matrix defined via the batch of sequences. The score equates to the normalized ratio of trace and determinant.

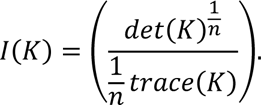

The score is valued between 0 and 1, where 1 is achieved once K forms essentially a diagonal matrix. If this is the case, this means the implicit features (defined via the kernel) are orthogonal to each other. On the other hand, 0 indicates that the implicit features defined via the kernel are very closely aligned to each other. Of course, this score depends on the kernel metric we use. The DPP method practically maximizes this metric under the models’ kernel in expectation.

### Clustering of ArM variants

The clustering shown in Fig. 5 was created using the t-SNE (t-distributed stochastic neighbour embedding)^54^ clustering methodology. For this analysis, we used the kernel matrix of the chemical descriptor model. This model is based on a Gaussian process with ARD (automatic relevance determination) kernel length scales. The t-SNE algorithm clusters the data based on a similarity metric that includes exponentiated negative Euclidean distances. This is very similar to our machine learning model, with the exception that instead of a pure exponential, we use the Matérn kernel. However, this should qualitatively lead to similar results. Hence, to generate the clustering, we took the chemical descriptors, scaled them with appropriate length scales, and used the scikit-learn implementation of the t-SNE algorithm to generate the clusters. We tested several values of complexity, and the plotted clusterings correspond to a value of 150, as it appeared to generate the most structured results.

### Subsampling analysis

To analyse the effect of data set size on the predictive ability of the model, we created 20 random subsamples of the original data set for each subsampling fraction (0.1 - 1 in intervals of 0.1). We then applied the previously described machine learning pipeline, starting with the feature selection. To analyse the performance of the models, we used them to predict the activity of all ArM variants that were tested in the exploitation round, and calculated the mean squared error of the predictions as well as the precision in predicting hits (i.e., ArM variants with a higher activity than the reference variant). Precision is defined as the percentage of true hits among predicted hits. To investigate the effect of the exploration round, we calculated the precision of a model that was trained on all data from the initial library and the exploration round. In the latter case, the precision is different from the experimentally determined hit rate as not all experimentally tested variants were predicted to be hits by the model used here.

### Data and code availability

The data and code will be made available upon publication of the manuscript.

## Acknowledgments

The authors thank Fadri Christoffel for synthesizing the gold cofactor. This work was created as part of the NCCR Catalysis and the NCCR Molecular Systems Engineering, both National Centres of Competence in Research funded by the Swiss National Science Foundation. R.T. acknowledges a grant from the Naito Foundation.

## Author contributions

T.V. and M.J. conceived the project. T.V. and G.S. developed the automated screening methods. T.V. and C.S. performed experiments. T.V. analysed screening results and NGS data. M.M. developed, applied, and analysed the machine learning pipeline. R.T. developed initial computational models. M.J., S.P. and T.R.W. supervised experimental work. A.K. supervised machine learning aspects. T.V., M.M. and M.J. wrote the manuscript with input from all authors.

