## Supplementary Information for "Enhanced Sequence-Activity Mapping and Evolution of Artificial Metalloenzymes by Active Learning"

#### Supplementary Figures

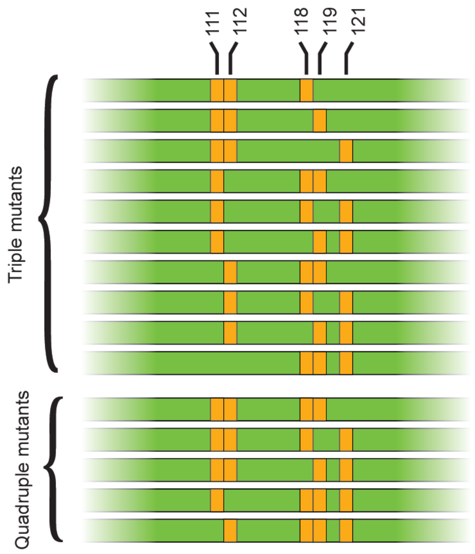


**Supplementary Fig. 1 | Illustration of the library design comprising both triple and quadruple mutant sub-libraries.** The orange rectangles indicate randomized positions in the Sav gene. When keeping two or one out of five positions constant, it is possible to create 10 sets of triple mutants and 5 sets of quadruple mutants. Note that the reference variant Sav S112F K121Q was selected as the parent of this library.

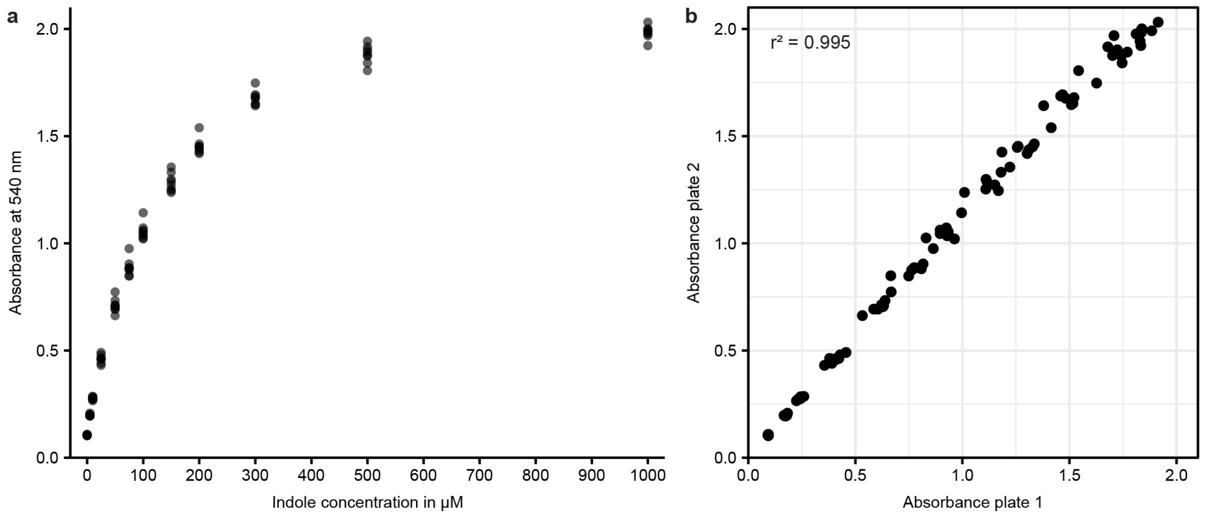


**Supplementary Fig. 2 | Validation of the automated Kovac's assay. a**, Indole standard curve measured using the automated Kovac's assay. Eight replicates were measured per concentration and all samples were in the same 96-well plate. Note that the indole concentrations observed in screenings of ArM mutants are below 100 µM. **b**, Reproducibility of indole measurements using the automated Kovac's assay. Two 96-well plates were filled with identical indole standards and subjected to the automated assay. The absorbance values of corresponding samples are plotted against each other and show a high correlation as determined by linear regression.

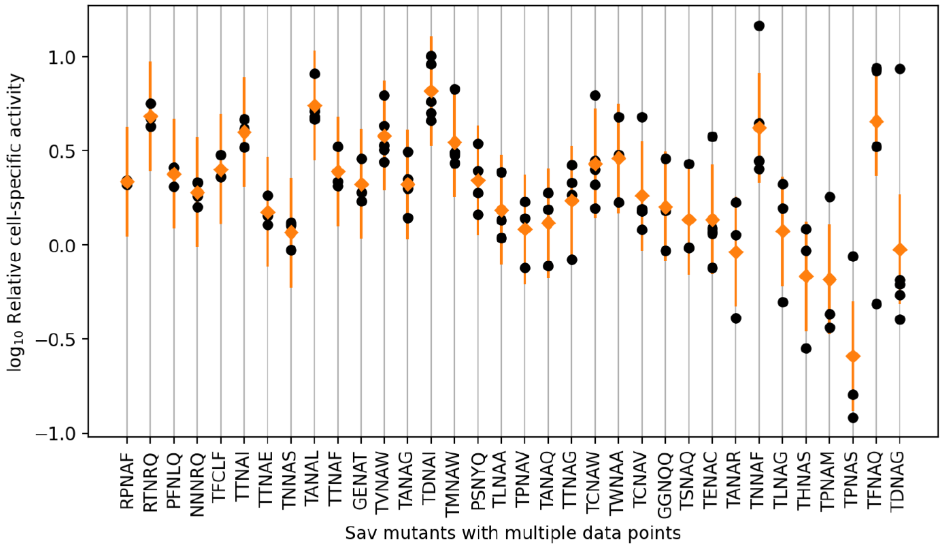


**Supplementary Fig. 3 | Reproducibility in the initial screening.** Points on the vertical lines correspond to variants that were found more than twice in the library. The variance between these replicates increases from left to right. The orange diamonds indicate the mean of the replicates and the orange lines display the two-fold standard deviation.

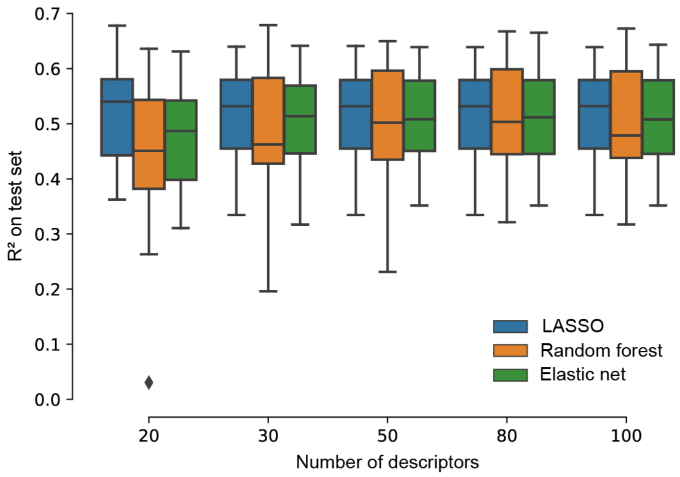


**Supplementary Fig. 4 | Benchmarking of the feature selection mechanism to create a subset for the final Bayesian model selection with automatic relevance detection.** We benchmarked three methods: two linear methods (LASSO and elastic net), and one non-linear method (random forest using its feature relevance score). LASSO generally performed best, particularly in selecting 20 features, likely due to its simplicity and robustness against noise. All methods were implemented using the scikit-learn toolkit. The benchmarking was done using 15-fold cross validation making sure no duplicates are shared between test and train.

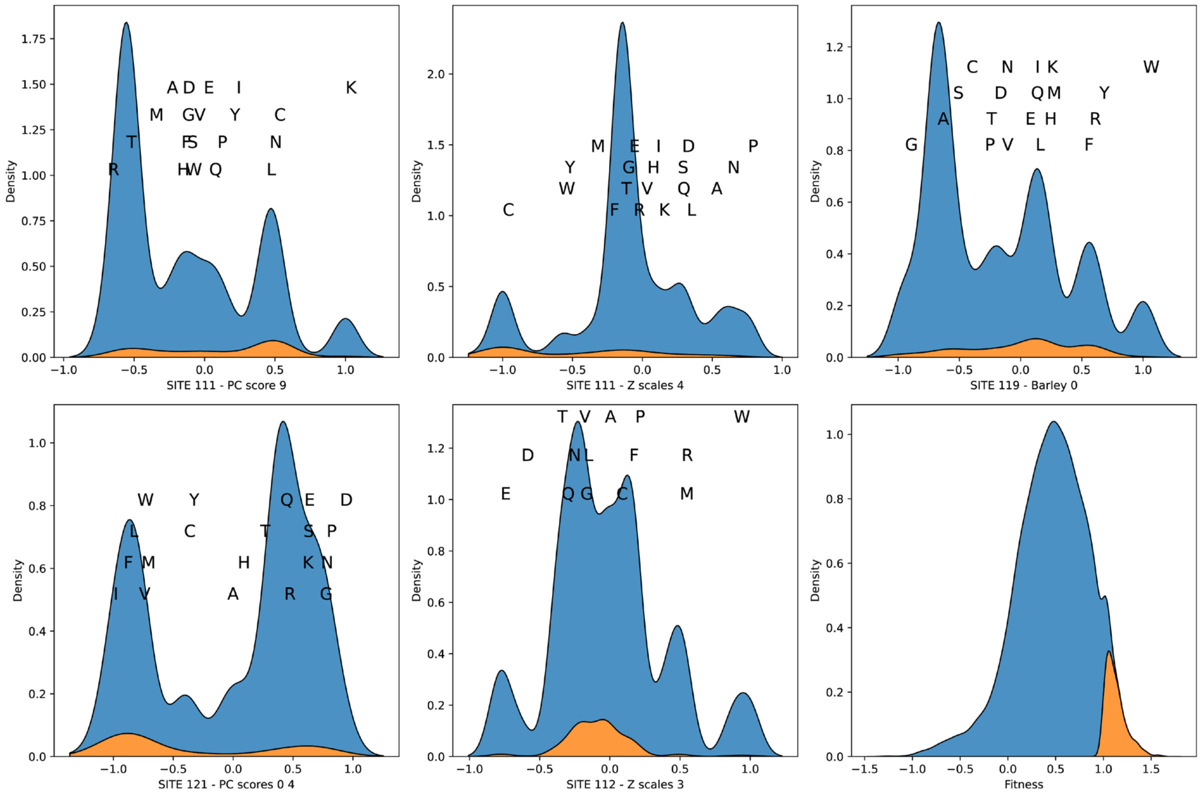


**Supplementary Fig. 5 | Distribution of the five most important features in the data set.** The blue area represents the background distribution, while the orange highlights the distribution of variants that are more active than the reference variant Sav TFNAQ. Notably, the orange distribution sometimes appears as a single peak and other times as bimodal, suggesting multiple ways of achieving high activity. The characters mark the value of the given feature that corresponds to the respective amino acid. The plots also demonstrate that different amino acids at specific sites can result in identical feature values. For example, as shown in the upper-left panel, amino acids D, G, F, and H all yield the same feature value. The bottom-right panel displays the distribution of activity values. The names of the features are given in the x-label.

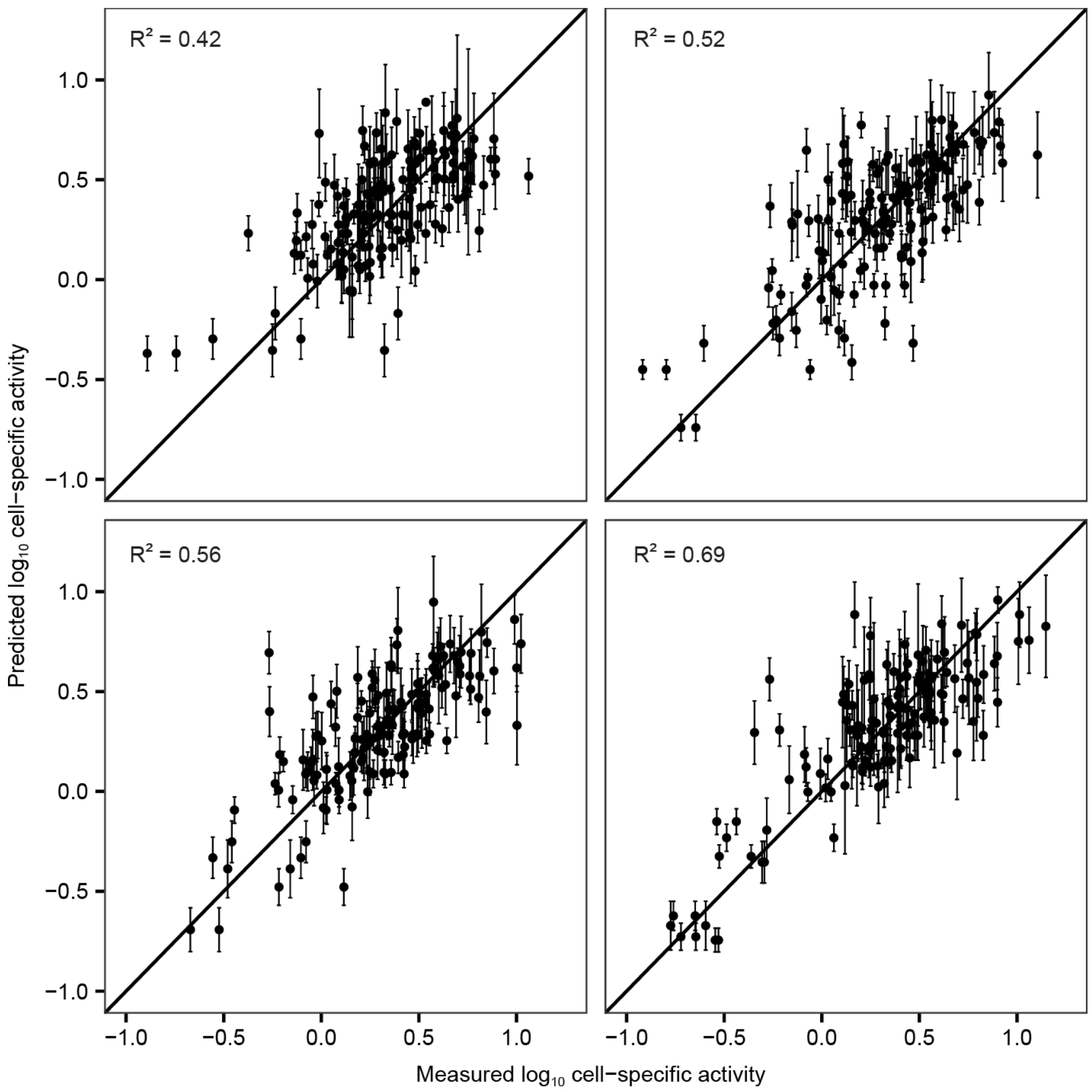


**Supplementary Fig. 6 | Exemplary cross-validation splits from the initial training phase.** The underlying model is the GP model using chemical descriptors with 20 features. The top left graph depicts the validation split with the lowest correlation, while the bottom right graph shows the split with the highest correlation. The other plots show splits with an intermediate correlation.

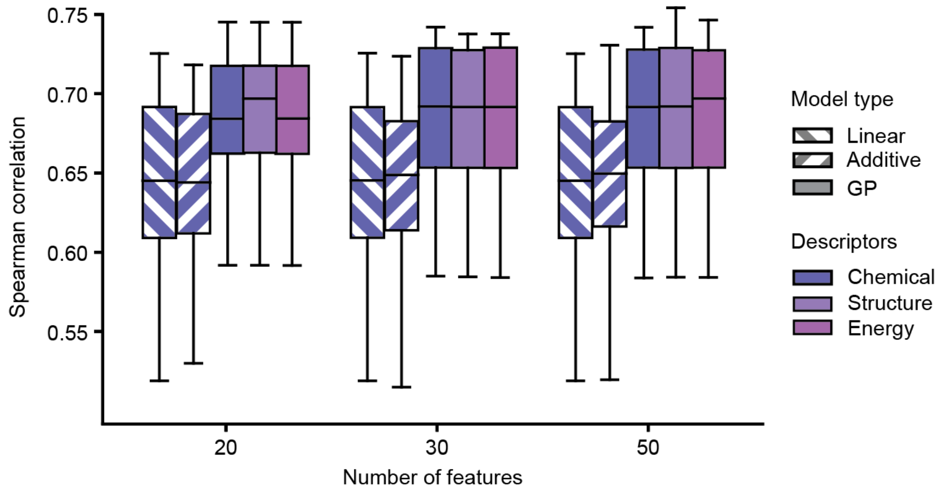


**Supplementary Fig. 7 | Spearman correlation (based on 15-fold cross-validation) of several models trained on the initial data set.** The influence of the number of features, model type, and descriptors was investigated.

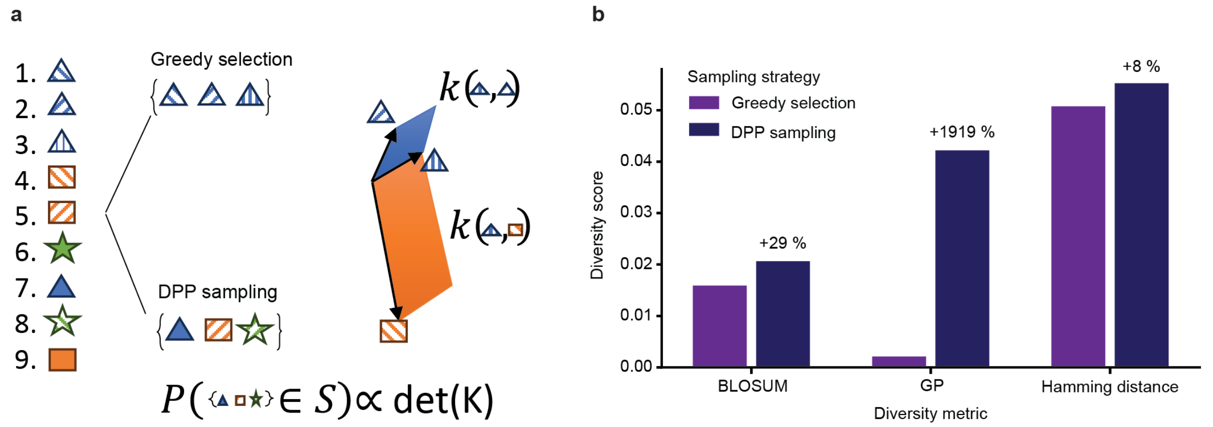


**Supplementary Fig. 8 | Diversification using determinantal point processes (DPP).** **a**, Illustration of DPP sampling. A ranked set of variants is depicted as pictograms. The same colour and shape indicate similar enzymes. Whereas greedy selection of top-ranked variants is likely to result in a set of highly similar variants, DPP sampling selects a more diverse batch (left). DPP sampling picks candidates based on the volume that their representations span in Euclidian space (right). More different variants span a larger volume, hence there is a larger probability of them being selected. An idealized batch would be such that all vectors are orthogonal to each other, forming a hypercube. **b**, Comparison of two sampling strategies in the exploitation round: Diversified selection based on DPP and greedy selection of the most active variants (predicted by the model using chemical descriptors). The diversity of the two sets of variants was compared using three metrics: BLOSUM90 substitution matrix, the model-based metric (GP), and the Hamming distance, sometimes also referred to as one-hot distance. Note that the different metrics are only expected to correlate weakly with each other. As the GP model was used to make the selection, the considerable increase in diversity according to this metric is expected.

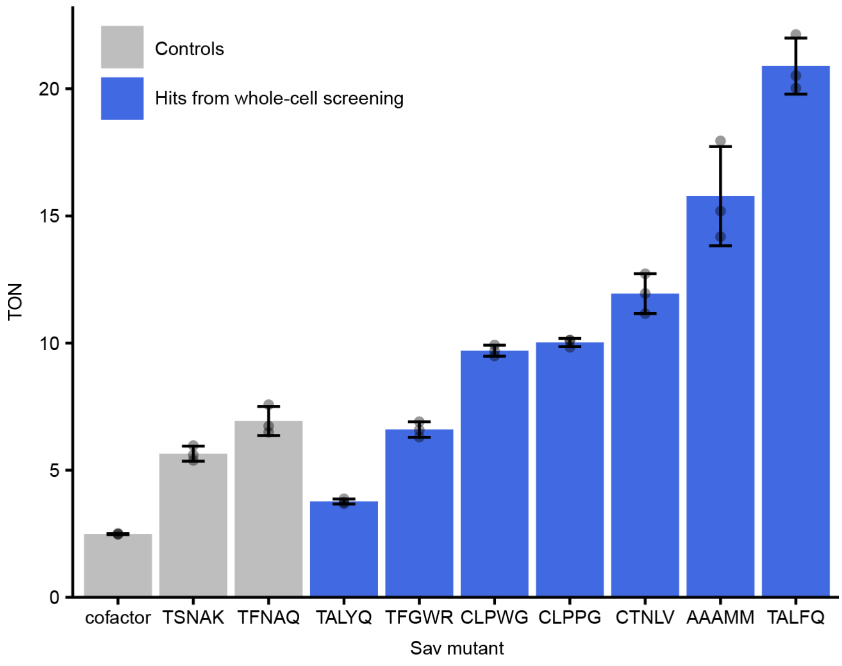


**Supplementary Fig. 9 | *In vitro* turnover number (TON) of ArM variants identified in the whole-cell screening.** The Sav variants with the highest cell-specific activity for hydroamination according to the validation experiment (Fig. 4d) were purified. Reactions were performed in triplicate with 10 µM Sav biotin-binding sites, 5 µM **(Biot-NHC)Au1** and 5 mM 2-ethynylaniline. For comparison, reactions were also performed using the free cofactor without Sav. After 24 h, the indole concentration was determined using the Kovac's assay. TSNAK is the wild-type variant, and TFNAQ refers to the best variant identified in a previous screening. Note that the TONs determined here are lower than in previous studies, possibly due to a less active batch of **(Biot-NHC)Au1**.

### Supplementary Tables

**Supplementary Table 1 | Most informative features identified by automatic relevance detection for Gaussian processes.** Most features are not interpretable, except for the Barley score which correlates with the size of the amino acid at position 119. When we overlay the distribution of the active vs. non-active variants, we discover that these features for active variants follow three different patterns. Optimal – meaning all good variants have this feature corresponding to one value, approximately bimodal – there are roughly two good values for these features, or strongly-bimodal – representing two equally good options for this feature.  A visual depiction of this analysis can be found in Supplementary Fig. 5.

| Feature | Site | Interpretation | Effect |
|---|---|---|---|
| PC 9 | 111 | Chemical | Approximately Bimodal |
| Z-score 4 | 111 | Chemical | Approximately Bimodal |
| Barley 0 | 119 | Chemical (size) | Tri-modal |
| PC 0 | 121 | Chemical | Strongly Bimodal |
| Z-score 3 | 121 | Chemical | Optimal |

**Supplementary Table 2 | Primers used for site-saturation mutagenesis at 20 Sav positions in proximity to the cofactor.** Mutations were introduced using NDT codons, which are highlighted in bold.

| Name | Sequence (5’ → 3’) |
|---|---|
| SSM_47_NDT_fwd | TGACCGGAACCTACGAGTCGGCC**NDT**GGCAACGCCGAGAGC |
| SSM_48_NDT_fwd | CCGGAACCTACGAGTCGGCCGTC**NDT**AACGCCGAGAGCCGC |
| SSM_49_NDT_fwd | GAACCTACGAGTCGGCCGTCGGC**NDT**GCCGAGAGCCGCTAC |
| SSM_86_NDT_fwd | CCTGGAAGAATAACTACCGCAAC**NDT**CACTCCGCGACCACG |
| SSM_87_NDT_fwd | GGAAGAATAACTACCGCAACGCC**NDT**TCCGCGACCACGTGG |
| SSM_88_NDT_fwd | AGAATAACTACCGCAACGCCCAC**NDT**GCGACCACGTGGAGC |
| SSM_110_NDT_fwd | GGCGAGGATCAACACCCAGTGGCTG**NDT**ACCTTTGGCACCACCGAG |
| SSM_111_NDT_fwd | GAGGATCAACACCCAGTGGCTGCTG**NDT**TTTGGCACCACCGAGG |
| SSM_112_NDT_fwd | GATCAACACCCAGTGGCTGCTGACC**NDT**GGCACCACCGAGGC |
| SSM_113_NDT_fwd | CAACACCCAGTGGCTGCTGACCTTT**NDT**ACCACCGAGGCCAACG |
| SSM_114_NDT_fwd | CACCCAGTGGCTGCTGACCTTTGGC**NDT**ACCGAGGCCAACGCC |
| SSM_115_NDT_fwd | CCAGTGGCTGCTGACCTTTGGCACC**NDT**GAGGCCAACGCCTGG |
| SSM_117_NDT_fwd | GCTGCTGACCTTTGGCACCACCGAG**NDT**AACGCCTGGCAATCCAC |
| SSM_118_NDT_fwd | GCTGACCTTTGGCACCACCGAGGCC**NDT**GCCTGGCAATCCACGC |
| SSM_119_NDT_fwd | GACCTTTGGCACCACCGAGGCCAAC**NDT**TGGCAATCCACGCTGGT |
| SSM_120_NDT_fwd | CTTTGGCACCACCGAGGCCAACGCC**NDT**CAATCCACGCTGGTCG |
| SSM_121_NDT_fwd | TGGCACCACCGAGGCCAACGCCTGG**NDT**TCCACGCTGGTCGG |
| SSM_122_NDT_fwd | CACCACCGAGGCCAACGCCTGGCAA**NDT**ACGCTGGTCGGCCAC |
| SSM_123_NDT_fwd | CACCGAGGCCAACGCCTGGCAATCC**NDT**CTGGTCGGCCACGACAC |
| SSM_124_NDT_fwd | CGAGGCCAACGCCTGGCAATCCACG**NDT**GTCGGCCACGACACCTT |
| kanR_rev | CAACGGGAAACGTCTTGCTC |
| kanR_fwd | ATTTAATCGCGGCCTAGAGC |
| SSM_47_rev | GGCCGACTCGTAGGTTC |
| SSM_48_rev | GACGGCCGACTCGTAGG |
| SSM_49_rev | GCCGACGGCCGACTC |
| SSM_86_rev | GTTGCGGTAGTTATTCTTCCAGG |
| SSM_87_rev | GGCGTTGCGGTAGTTATTC |
| SSM_88_rev | GTGGGCGTTGCGGTAGTTAT |
| SSM_110_rev | CAGCCACTGGGTGTTGATC |
| SSM_111_rev | CAGCAGCCACTGGGTGTTG |
| SSM_112_rev | GGTCAGCAGCCACTGGG |
| SSM_113_rev | AAAGGTCAGCAGCCACTGG |
| SSM_114_rev | GCCAAAGGTCAGCAGCCAC |
| SSM_115_rev | GGTGCCAAAGGTCAGCAG |
| SSM_117_rev | CTCGGTGGTGCCAAAGGTC |

| Name | Sequence (5' → 3') |
|---|---|
| SSM_118_rev | GGCCTCGGTGGTGC |
| SSM_119_rev | GTTGGCCTCGGTGGTG |
| SSM_120_rev | GGCGTTGGCCTCGGTG |
| SSM_121_rev | CCAGGCGTTGGCCTCG |
| SSM_122_rev | TTGCCAGGCGTTGGCC |
| SSM_123_rev | GGATTGCCAGGCGTTGG |
| SSM_124_rev | CGTGGATTGCCAGGCGTTG |

**Supplementary Table 3 | Primers used to generate libraries of double, triple, quadruple and quintuple mutants.** Degenerate codons are highlighted in bold.

| Name | Sequence (5' → 3') |
|---|---|
| Sav_FQ_fwd | GGCACCACCGAGGCCAACGCCTGGCAATCCACGCTGGTCGGC |
| Sav_118X_fwd | GGCACCACCGAGGCC**NNK**GCCTGGCAATCCACGCTGGTCGGC |
| Sav_119X_fwd | GGCACCACCGAGGCCAAC**NNK**TGGCAATCCACGCTGGTCGGC |
| Sav_121X_fwd | GGCACCACCGAGGCCAACGCCTGG**NNK**TCCACGCTGGTCGGC |
| Sav_118X_119X_fwd | GGCACCACCGAGGCC**NNKNNK**TGGCAATCCACGCTGGTCGGC |
| Sav_118X_121X_fwd | GGCACCACCGAGGCC**NNK**GCCTGG**NNK**TCCACGCTGGTCGGC |
| Sav_119X_121X_fwd | GGCACCACCGAGGCCAAC**NNK**TGG**NNK**TCCACGCTGGTCGGC |
| Sav_118X_119X_121X_fwd | GGCACCACCGAGGCC**NNKNNK**TGG**NNK**TCCACGCTGGTCGGC |
| Sav_FQ_rev | GGCCTCGGTGGTGCCAAAGGTCAGCAGCCACTGGGTG |
| Sav_111X_rev | GGCCTCGGTGGTGCCAAA**MNN**CAGCAGCCACTGGGTG |
| Sav_112X_rev | GGCCTCGGTGGTGCC**MNN**GGTCAGCAGCCACTGGGTG |
| Sav_111X_112X_rev | GGCCTCGGTGGTGCC**MNNMNN**CAGCAGCCACTGGGTG |
| kanR_fwd | ATTTAATCGCGGCCTAGAGC |
| kanR_rev | CAACGGGAAACGTCTTGCTC |

**Supplementary Table 4 | Primer combinations used to generate fragments with mutations at positions 111 and 112 or positions 118, 119, and 121.** The primer sequences can be found in Supplementary Table 3.

| | Forward primer(s) | Reverse primer(s) |
|---|---|---|
| PCR 1 | Sav_FQ_fwd | kanR_rev |
| PCR 2 | Sav_118X_fwd<br>Sav_119X_fwd<br>Sav_121X_fwd | kanR_rev |
| PCR 3 | Sav_118X_119X_fwd<br>Sav_118X_121X_fwd<br>Sav_119X_121X_fwd | kanR_rev |
| PCR 4 | Sav_118X_119X_121X_fwd | kanR_rev |
| PCR 5 | kanR_fwd | Sav_FQ_rev |
| PCR 6 | kanR_fwd | Sav_111X_rev<br>Sav_112X_rev |
| PCR 7 | kanR_fwd | Sav_111X_112X_rev |

**Supplementary Table 5 | Assembly scheme for fragments from Supplementary Table 4.**

| Assembly reaction | Type of mutants | Fragment A | Fragment B | Theoretical diversity |
|---|---|---|---|---|
| 1 | Double | PCR 7 | PCR 1 | 400 |
| 2 | Double | PCR 6 | PCR 2 | 2,400 |
| 3 | Double | PCR 5 | PCR 3 | 1,200 |
| 4 | Triple | PCR 7 | PCR 2 | 24,000 |
| 5 | Triple | PCR 6 | PCR 3 | 48,000 |
| 6 | Triple | PCR 5 | PCR 4 | 8,000 |
| 7 | Quadruple | PCR 7 | PCR 3 | 480,000 |
| 8 | Quadruple | PCR 6 | PCR 4 | 320,000 |
| 9 | Quintuple | PCR 7 | PCR 4 | 3,200,000 |

**Supplementary Table 6 | Primers used to create libraries of specific variants suggested by the machine learning model.** The positions at which the oligo pools contained diverse nucleotides are represented by Ns and are highlighted in bold.

| Description | Sequence (5' → 3') |
|---|---|
| Oligo pool | GAGGATCAACACCCAGTGGCTGCTG**NNNNNN**GGCACCACCGAGGCC**NNNNNN**TGG**NNN**TCCACGCTGGTCGGCCACG |
| kanR_rev_B | CAACGGGAAACGTCTTGCTCTAGGCC |
| ML_amp_fwd | GAGGATCAACACCCAGTGGC |
| kanR_rev | CAACGGGAAACGTCTTGCTC |
| kanR_fwd | ATTTAATCGCGGCCTAGAGC |
| SSM_111_rev | CAGCAGCCACTGGGTGTTG |

**Supplementary Table 7 | PCRs conducted to clone the first library of ML-designed Sav variants.** For PCR 1, 15 cycles were performed with the initial primer combination, at which point the other primers were added for 20 additional cycles to amplify the product further. The primer sequences can be found in Supplementary Table 6.

| | Forward primer | Concentration | Reverse primer | Concentration | Cycles |
|---|---|---|---|---|---|
| PCR 1 | Oligo pool | 2 nM | kanR_rev_B | 500 nM | 15 |
| | ML_amp_fwd | 500 nM | kanR_rev | 500 nM | 20 |
| PCR 2 | kanR_fwd | 500 nM | SSM_111_rev | 500 nM | 30 |

**Supplementary Table 8 | PCRs conducted to clone the second library of ML-designed Sav variants.** The primer sequences can be found in Supplementary Table 6.

| | Forward primer | Concentration | Reverse primer | Concentration | Cycles |
|---|---|---|---|---|---|
| PCR 1 | Oligo pool | 40 nM | kanR_rev_B | 500 nM | 35 |
| PCR 2 | kanR_fwd | 500 nM | SSM_111_rev | 500 nM | 30 |

**Supplementary Table 9 | Primers for well-specific barcoding of Sav mutants.** Barcodes are highlighted in bold.

| Description | Sequence (5' → 3') |
|---|---|
| Sav_seq_rowA_fwd | GAGACCTTGTGTCGATGG**GGAGAA**GACCGGAACCTACGAGTCGGCC |
| Sav_seq_rowB_fwd | GAGACCTTGTGTCGATGG**CTGGAA**GACCGGAACCTACGAGTCGGCC |
| Sav_seq_rowC_fwd | GAGACCTTGTGTCGATGG**TCCGAA**GACCGGAACCTACGAGTCGGCC |
| Sav_seq_rowD_fwd | GAGACCTTGTGTCGATGG**ACAGTG**GACCGGAACCTACGAGTCGGCC |
| Sav_seq_rowE_fwd | GAGACCTTGTGTCGATGG**GTCTAG**GACCGGAACCTACGAGTCGGCC |
| Sav_seq_rowF_fwd | GAGACCTTGTGTCGATGG**CTCTTC**GACCGGAACCTACGAGTCGGCC |
| Sav_seq_rowG_fwd | GAGACCTTGTGTCGATGG**TATCGC**GACCGGAACCTACGAGTCGGCC |
| Sav_seq_rowH_fwd | GAGACCTTGTGTCGATGG**AGTAGG**GACCGGAACCTACGAGTCGGCC |
| Sav_seq_col1_rev | GTCACACGTAGCATGTGG**CATAGG**CCTTGGTGAAGGTGTCGTGGCC |
| Sav_seq_col2_rev | GTCACACGTAGCATGTGG**TCCGTT**CCTTGGTGAAGGTGTCGTGGCC |
| Sav_seq_col3_rev | GTCACACGTAGCATGTGG**CATGCA**CCTTGGTGAAGGTGTCGTGGCC |
| Sav_seq_col4_rev | GTCACACGTAGCATGTGG**TTGTGG**CCTTGGTGAAGGTGTCGTGGCC |
| Sav_seq_col5_rev | GTCACACGTAGCATGTGG**TTGCCT**CCTTGGTGAAGGTGTCGTGGCC |
| Sav_seq_col6_rev | GTCACACGTAGCATGTGG**TTGGTC**CCTTGGTGAAGGTGTCGTGGCC |
| Sav_seq_col7_rev | GTCACACGTAGCATGTGG**CTCTTG**CCTTGGTGAAGGTGTCGTGGCC |
| Sav_seq_col8_rev | GTCACACGTAGCATGTGG**GGACAA**CCTTGGTGAAGGTGTCGTGGCC |
| Sav_seq_col9_rev | GTCACACGTAGCATGTGG**GACTTC**CCTTGGTGAAGGTGTCGTGGCC |
| Sav_seq_col10_rev | GTCACACGTAGCATGTGG**CCAATC**CCTTGGTGAAGGTGTCGTGGCC |
| Sav_seq_col11_rev | GTCACACGTAGCATGTGG**CAACGA**CCTTGGTGAAGGTGTCGTGGCC |
| Sav_seq_col12_rev | GTCACACGTAGCATGTGG**AAGGCT**CCTTGGTGAAGGTGTCGTGGCC |

**Supplementary Table 10 | Primers for plate-specific barcoding of Sav mutants.** Barcodes are highlighted in bold.

| Description | Sequence (5' → 3') |
|---|---|
| Sav_seq_ext1_fwd | CAAGCAGAAGACGGCATACGAGATGTGACTGGAGTTCAGACGTGTGCTCTTCCGATCT**GGTAC**GAGACCTTGTGTCGATGG |
| Sav_seq_ext2_fwd | CAAGCAGAAGACGGCATACGAGATGTGACTGGAGTTCAGACGTGTGCTCTTCCGATCTC**AACAC**GAGACCTTGTGTCGATGG |
| Sav_seq_ext3_fwd | CAAGCAGAAGACGGCATACGAGATGTGACTGGAGTTCAGACGTGTGCTCTTCCGATCTAT**CGGTT**GAGACCTTGTGTCGATGG |
| Sav_seq_ext4_fwd | CAAGCAGAAGACGGCATACGAGATGTGACTGGAGTTCAGACGTGTGCTCTTCCGATCTTCG**GTCAA**GAGACCTTGTGTCGATGG |
| Sav_seq_ext5_fwd | CAAGCAGAAGACGGCATACGAGATGTGACTGGAGTTCAGACGTGTGCTCTTCCGATCTATCG**AAGCG**GAGACCTTGTGTCGATGG |
| Sav_seq_ext6_fwd | CAAGCAGAAGACGGCATACGAGATGTGACTGGAGTTCAGACGTGTGCTCTTCCGATCTG**CCACA**GAGACCTTGTGTCGATGG |
| Sav_seq_ext1_rev | AATGATACGGCGACCACCGAGATCTACACTCTTTCCCTACACGACGCTCTTCCGATCT**AGGAA**GTCACACGTAGCATGTGG |
| Sav_seq_ext2_rev | AATGATACGGCGACCACCGAGATCTACACTCTTTCCCTACACGACGCTCTTCCGATCTG**AGTGG**GTCACACGTAGCATGTGG |
| Sav_seq_ext3_rev | AATGATACGGCGACCACCGAGATCTACACTCTTTCCCTACACGACGCTCTTCCGATCTCC**ACGTC**GTCACACGTAGCATGTGG |
| Sav_seq_ext4_rev | AATGATACGGCGACCACCGAGATCTACACTCTTTCCCTACACGACGCTCTTCCGATCTTTC**TCAGC**GTCACACGTAGCATGTGG |
| Sav_seq_ext5_rev | AATGATACGGCGACCACCGAGATCTACACTCTTTCCCTACACGACGCTCTTCCGATCTCAAG**CTAGC**GTCACACGTAGCATGTGG |
| Sav_seq_ext6_rev | AATGATACGGCGACCACCGAGATCTACACTCTTTCCCTACACGACGCTCTTCCGATCTT**GCTTA**GTCACACGTAGCATGTGG |

**Supplementary Table 11 | Features used for the chemical descriptors.**

| Feature group name | Number of features | Description | Source |
|---|---|---|---|
| Z-scales | 8 | PCA score | [16] |
| VHSE | 5 | PCA score | [17] |
| Barley | 2 | PCA score | [18] |
| PCscores | 11 | PCA score | [55] |

**Supplementary Table 12 | Geometric features calculated for dimeric Sav.** Notice that these are raw features. Subsequently, we discarded features that did not vary across the 3.2 million structures, leaving us with 682 features. Solvent-accessible surface area is abbreviated SASA.

| Feature group name | Number of features | Description | Source |
|---|---|---|---|
| SASA 1.4 | 238 | Rolling ball alg. | Biotite[69] |
| SASA 1.4 H | 238 | Rolling ball alg. | Biotite |
| SASA 3.5 | 238 | Rolling ball alg. | Biotite |
| SASA 3.5 H | 238 | Rolling ball alg. | Biotite |
| SASA 5.5 | 238 | Rolling ball alg. | Biotite |
| SASA 5.5 H | 238 | Rolling ball alg. | Biotite |
| Partial charges | 238 | Gasteiger method | Biotite |
| Centre of mass | 238*3 | Min ball method | Minball |
| Radius | 238*3 | Min ball method | Minball |
| Dihedrals | 238 | | Biotite |
| Hydrogen bonds per residue | 238 | [76] | Biotite |

**Supplementary Table 13 | Ref2015 score components from the Rosetta Suite[77].** These scores were calculated for each approximate structure.

| Feature name | Description |
|---|---|
| fa_atr | Lennard-Jones attractive between atoms in different residues |
| fa_rep | Lennard-Jones repulsive between atoms in different residues |
| fa_sol | Lazaridis-Karplus solvation energy |
| fa_intra_sol_xover4 | Intra-residue Lazaridis-Karplus solvation energy |
| lk_ball_wtd | Asymmetric solvation energy |
| fa_intra_rep | Lennard-Jones repulsive between atoms in the same residue |
| fa_elec | Coulombic electrostatic potential with a distance-dependent dielectric |
| pro_close | Proline ring closure energy and energy of psi angle of preceding residue |
| hbond_sr_bb | Backbone-backbone hbonds close in primary sequence |
| hbond_lr_bb | Backbone-backbone hbonds distant in primary sequence |
| hbond_bb_sc | Sidechain-backbone hydrogen bond energy |
| hbond_sc | Sidechain-sidechain hydrogen bond energy |
| dslf_fa13 | Disulfide geometry potential |
| rama_prepro | Ramachandran preferences (with separate lookup tables for pre-proline positions and other positions) |
| omega | Omega dihedral in the backbone. A Harmonic constraint on planarity with standard deviation of ~6 deg. |
| p_aa_pp | Probability of amino acid, given torsion values for phi and psi |
| fa_dun | Internal energy of sidechain rotamers as derived from Dunbrack's statistics |
| yhh_planarity | A special torsional potential to keep the tyrosine hydroxyl in the plane of the aromatic ring |
| ref | Reference energy for each amino acid. Balances internal energy of amino acid terms. Plays a role in design. |

**Supplementary Table 14 | Allocation of the experimental budget in the exploration round.**

| Group | Number of Sav variants |
|---|---|
| Informative | 504 |
| Conservative prediction | 72 |
| Balanced prediction | 72 |
| Optimistic prediction | 72 |

**Supplementary Table 15 | Allocation of the experimental budget in the exploitation round.**

| Group | Model | Number of Sav variants |
|---|---|---|
| Conservative prediction | Chemical | 120 |
| Balanced prediction | Chemical | 120 |
| Conservative prediction | Geometric | 120 |
| Balanced prediction | Geometric | 120 |
| Conservative prediction | Energy-based | 120 |
| Balanced prediction | Energy-based | 120 |
